# Structural framework for the understanding spectroscopic and functional signatures of the cyanobacterial Orange Carotenoid Protein families

**DOI:** 10.1101/2023.08.20.554024

**Authors:** Nikolai N. Sluchanko, Eugene G. Maksimov, Yury B. Slonimskiy, Larisa A. Varfolomeeva, Antonina Y. Bukhanko, Nikita A. Egorkin, Georgy V. Tsoraev, Maria G. Khrenova, Baosheng Ge, Song Qin, Konstantin M. Boyko, Vladimir O. Popov

## Abstract

The Orange Carotenoid Protein (OCP) is a unique photoreceptor crucial for cyanobacterial photoprotection. Best studied *Synechocystis* sp. PCC 6803 OCP belongs to the large OCP1 family. Downregulated by the Fluorescence Recovery Protein (FRP) in low-light, high-light-activated OCP1 binds to the phycobilisomes and performs non-photochemical quenching. Recently discovered families OCP2 and OCP3 remain structurally and functionally underexplored, and no systematic comparative studies have ever been conducted. Here we present two first crystal structures of OCP2 from morphoecophysiologically different cyanobacteria and provide their comprehensive structural, spectroscopic and functional comparison with OCP1, the recently described OCP3 and all-OCP ancestor. Structures enable correlation of spectroscopic signatures with the effective number of hydrogen and discovered here *chalcogen* bonds anchoring the ketocarotenoid in OCP and rationalize the observed differences in OCP/FRP and OCP/phycobilisome functional interactions. These data are expected to foster OCP research and applications in optogenetics, targeted carotenoid delivery and cyanobacterial biomass engineering.

## Introduction

High light conditions threaten to photodamage the photosynthetic apparatus and require variable mechanisms of photoprotection. Cyanobacteria have evolved a unique tolerance mechanism based on the Orange Carotenoid Protein (OCP), a 35-kDa water-soluble photoactive carotenoprotein [1–5]. OCP has a modular organization [6]: its α-helical N-terminal domain (NTD) and the α/β-folded C-terminal domain (CTD) are connected by the long interdomain linker and share a single ketocarotenoid molecule: 3-hydroxyechinenone, echinenone (ECH) or canthaxanthin (CAN) [7,8]. Hydroxycarotenoid binding does not make OCP photoactive [9]. In the dark-adapted orange form, OCP^O^, the carotenoid keto group is coordinated by H-bonds with the side chains of the conserved Tyr201 and Trp288 residues (*Synechocystis* sp. PCC 6803 numbering) in the CTD [7,8,10]. The OCP^O^ form is stabilized by the NTD/CTD interactions at the major interdomain groove, including the salt bridge between Arg155 in the NTD and Glu244 in the CTD [8,11], and by the N-terminal extension (NTE) attached to the CTD [7,8,12–14]. Intense visible light (400-500 nm) triggers OCP transformation into the red form, OCP^R^, which binds to the phycobilisomes (PBS) and dissipates the excess absorbed light energy as heat [4,5,15].

The OCP^O^-OCP^R^ (O-R for simplicity) transition is a multistep, reversible process involving the disruption of the H-bonds with Tyr201/Trp288, carotenoid translocation 12 A towards the NTD to be fully accommodated in this domain, NTE detachment from its docking site on the CTD, and domain separation [16–21]. Mutation of Tyr/Trp residues (e.g., W288A or Y201A/W288A) yields an OCP^R^ mimic with a pronounced tendency to dimerization and persistent PBS fluorescence quenching ability [10,20]. The OCP^R^ form recruits the dimeric Fluorescence Recovery Protein (FRP), which facilitates the OCP domain re-association and accelerates the R-O transition [10,22–24], completely inactivating OCP in low light. The main FRP-binding site is located on the CTD region covered by the NTE in OCP^O^ [13,25], while the secondary sites on the NTD help FRP to re-associate the OCP domains [24] and recover PBS fluorescence [26]. The recently obtained cryo-EM structure of the PBS-OCP complex reveals the OCP-binding site within the PBS core [27].

Phylogenetic studies have found that at least three distinct OCP families exist – OCP1, OCP2 and OCP3 (initially named OCPX) [28]. The OCP1 genes often co-occur with FRP genes, sometimes co-occur with either OCP2 or OCP3 paralogous genes, whereas OCP2 and OCP3 genes are found regardless of FRP genes, as OCP2– or OCP3-only genomes do not usually encode FRP [28,29]. The canonical FRP-regulated photoprotection mechanism concerns only the largest OCP1 family, while OCP2 and OCP3 families remain largely underexplored. The heterogeneous OCP3 group apparently combines at least three subfamilies (OCP3a, OCP3b, OCP3c), with OCP3a from *Gloeobacteria* – the basal cyanobacteria group – being the most primitive and closest to the common OCP ancestor [30]. OCP3 representatives are studied for *Scytonema hofmanni* PCC 7110 [29], *Gloeobacter kilaueensis* JS1 [30] and *Nostoc flagelliforme* [31], with crystal structures for the latter two representatives reported in the last 2 years [30,31]. By contrast, OCP2 characterization [28,29,32] is lagging behind and concerned only on a single representative from *Tolypothrix* sp. PCC 7601 (encodes OCP1, OCP2 and FRP). Comparative analysis revealed that, while the constitutively expressed TolyOCP1 is canonically regulated by TolyFRP, the latter does not affect the photocycle of TolyOCP2, which in turn is expressed only in high light [28]. The available data were limited to CAN-functionalized, C-terminally His-tagged TolyOCP1 and TolyOCP2 [28,32]. While CAN is native to *Tolypothrix*, the type of the functionalizing carotenoid significantly influences properties of a given OCP [30,33,34]. The His-tags, especially at the C terminus, can strongly affect properties of OCP [33] and the C-terminal His-tag is partly seen in the TolyOCP1 structure (PDB ID: 6PQ1 [32]), where it forms nonnative stabilizing interactions with the CTD. No OCP2 structure could be obtained.

The scarce published data described OCP2 as a rather primitive, fast photoswitching OCP not regulated by FRP and less efficiently quenching PBS [28,29]. However, the rational explanations of all accumulated observations required structural data, which so far were missing only for the OCP2 family (Table 1). An OCP2-specific Gln155 residue in place of the highly conserved Arg155, involved in NTD-CTD and OCP-PBS interactions [8,11], left further intrigue [28]. Last but not least, there is growing demand in constructing various sensory, optogenetic devices and carotenoid carriers based on OCP with a controlled light response [35–38]. In addition, engineered OCP can accelerate recovery from photoprotection and potentially improve cyanobacterial biomass production [39,40]. Given the presumed different functional characteristics of OCP superfamily members, the progress in these directions could have been appreciably invigorated by a portfolio of OCP structures, but it still remained incomplete.

**Table 1.** Representative crystal structures of OCP from different families.

| PDB ID | Clade | Species | Carotenoid | Crystallographic oligomer | Largest intermonomer buried surface area (% of the total molecular SASA) | Ref. |
| --- | --- | --- | --- | --- | --- | --- |
| 5UI2 | OCP1 | Limnospira (Arthrospira) maxima | hECH | dimer | 1186.1 Å<sup>2</sup> (7.7%) | [7] |
| 3MG1 | OCP1 | Synechocystis sp. PCC 6803 | ECH | dimer | 1095.8 Å<sup>2</sup> (7.6%) | [8] |
| 5HGR | OCP1 | Anabaena (Nostoc) sp. PCC 7120 | CAN | dimer | 944.1 Å<sup>2</sup> (6.3%) | [82] |
| 6PQ1 | OCP1 | Tolypothrix sp. PCC 7601 | CAN | dimer | 856.9 Å<sup>2</sup> (5.5%) | [32] |
| 7EKR | OCP1 | Gloeocapsa sp. PCC 7513 | CAN | dimer | 932.5 Å<sup>2</sup> (6.2%) | [83] |
| 7QD2 | OCP1 | Planktothrix sp. PCC 7805 | CAN | dimer | 1083.7 Å<sup>2</sup> (7.5%) | [33] |
| 8A0H | OCP3 (OCPX) | Gloeobacter kilaueensis JS1 | ECH | monomer | 768.6 Å<sup>2</sup> (5.1%) | [30] |
| 7YTH | OCP3 (OCPX1) | Nostoc flagelliforme CCNUN1 | carotenoid not modeled due to incomplete electron density | monomer (oligomer in ASU) | 779.3 Å<sup>2</sup> (5.2%) | [31] |
| 7YTF | OCP3 (OCPX2) | Nostoc flagelliforme CCNUN1 | CAN | dimer* | 1290.9 Å<sup>2</sup> (8.8%) | [31] |
| 8PYH | OCP2 | Crinalium epipsammum PCC 9333 | ECH | dimer* | 860.7 Å<sup>2</sup> (5.6%) | this work |
| 8PZK | OCP2 | Gloeocapsa sp. PCC 7428 | ECH | dimer* | 1072.8 Å<sup>2</sup> (7.3%) | this work |

\*Crystallographic dimers significantly different from the typical crystallographic dimer of OCP1.

To fill these gaps, here we focus on two OCP2 members from morphoecophysiologically different cyanobacteria, describe their crystal structures and provide detailed structural, spectroscopic and functional comparison with OCP1, the recently described OCP3 [30,31] and the all-OCP ancestor (AncOCPall) [41]. We thus obtain a missing piece of the puzzle to propose a comprehensive structural framework for various analyses of the OCP superfamily, including properties of practical interest for nano– and biotechnology.

## Results

### OCP2 is the smallest and undercharacterized OCP family

The OCP2 family described in 2017 is the smallest and evolutionarily intermediate between OCP1 and OCP3 [28,29,41], having just one representative studied [28,29,32]. To expand knowledge of this least characterized family, for structural and functional OCP2 characterization we searched for cyanobacteria with markedly different ecological/morphological parameters. While revisiting the phylogenetic tree based on the recently published dataset [29] (Fig. 1A), we picked OCP2 representatives from *Crinalium epipsammum* PCC 9333 (CrinOCP2) (https://www.kegg.jp/kegg-bin/show_organism?org=cep) and *Gloeocapsa* sp. PCC 7428 (GcapOCP2) (https://www.kegg.jp/kegg-bin/show_organism?org=glp), in which the corresponding OCP2 gene is the sole OCP in the genome, and hence its functional properties may be most relevant to the physiology of the particular cyanobacterium. No FRP gene is found in either of these species. Both microorganisms contain genetic chassis for the biosynthesis of the ketocarotenoid echinenone (ECH) (https://www.kegg.jp/pathway/cep00906 and https://www.kegg.jp/pathway/glp00906). At least for *Crinalium* it was documented that, together with β-carotene, ECH is the major carotenoid [42]. This is a filamentous non-heterocyst and non-motile cyanobacterium highly resistant to drought, known for forming crusts in coastal sand dunes [43]. OCP-related proteins are known to be crucial for adaptation of terrestrial cyanobacteria to desiccation [44]. In turn, *Gloeocapsa* sp. PCC 7428 is a unicellular cyanobacterium found in moderate hot springs, closely related to heterocystous cyanobacteria [45]. These considerations made the formation of OCP2-echinenone complexes under native environment most probable and their reconstitution in available expression systems justified.

**Fig. 1.**
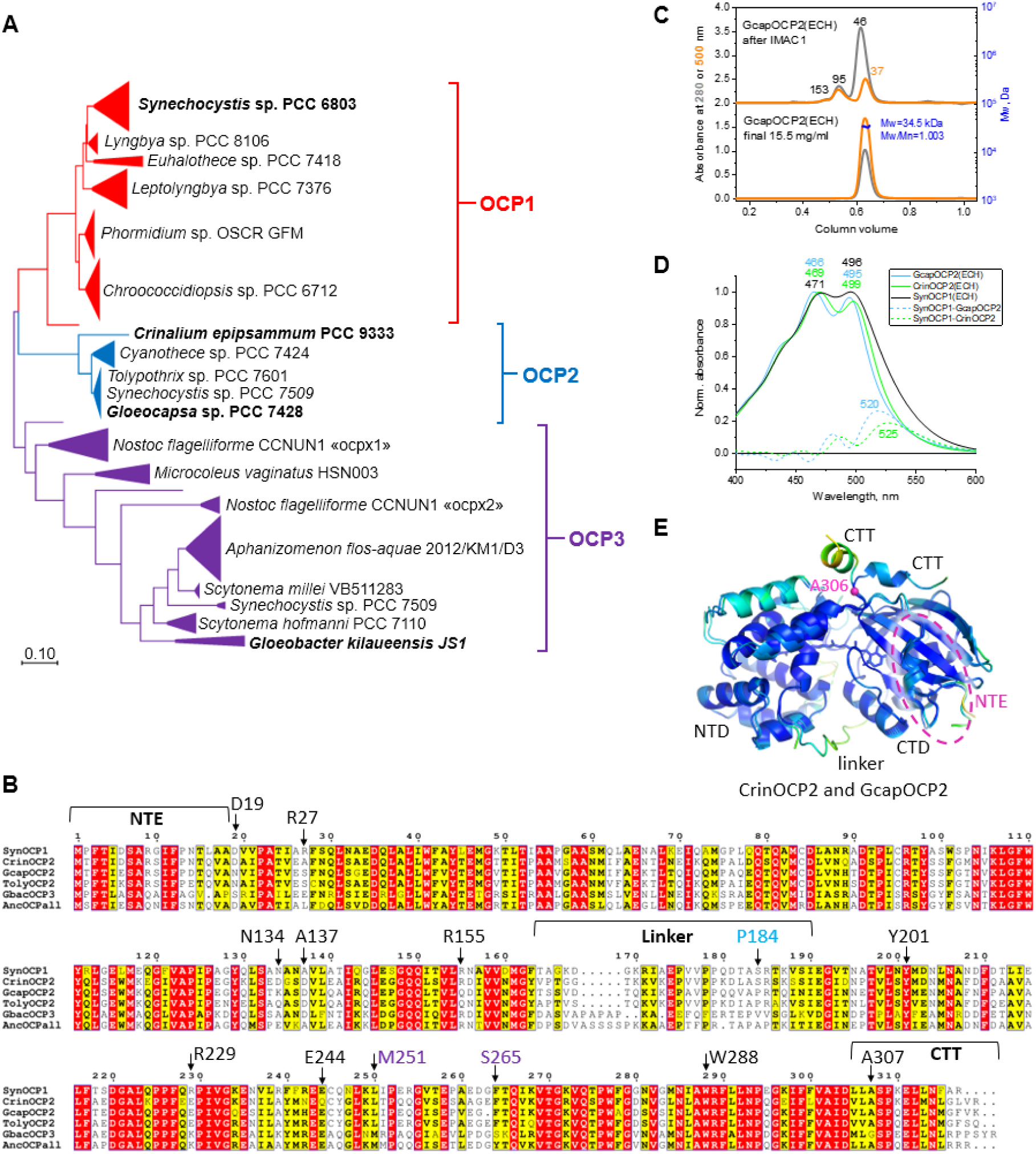
Selection and characterization of OCP2 members. **A**. Phylogenetic analysis of 150 OCP1, OCP2 and OCP3 sequences showing the location of the paralogs studied in this work (bold font). The tree with the highest log likelihood is presented and shown to scale, branch lengths reflect the number of substitutions per site (gaps were eliminated). **B**. Alignment of OCP sequences from *Synechocystis* sp. PCC 6803 (SynOCP1), *Crinalium epipsammum* PCC 9333 (CrinOCP2), *Gloeocapsa* sp. PCC 7428 (GcapOCP2), *Tolypothrix* sp. PCC 7601 (TolyOCP2), *Gloeobacter kilaueensis* JS1 (GbacOCP3) and the predicted ancestor sequence of all OCPs (AncOCPall; taken from [41]) built using ESPript (https://espript.ibcp.fr). Identical residues are highlighted by red, similar – by yellow. Similarity coloring scheme used the global similarity score of 0.7. Positions discussed in the text are marked (and color-coded for GbacOCP3 and GcapOCP2). **C**. SEC of GcapOCP2 expressed in ECH-producing *E.coli* cells, immediately after IMAC1 and after complete purification. The elution profiles are shifted along the Y axis for clarity. The apparent Mw for the peaks are indicated as determined from column calibration. The absolute Mw of the final GcapOCP2(ECH) preparation determined by SEC-MALS is shown for the final preparation along with the average Mw across the peak and the polydispersity index (Mw/Mn). **D**. Normalized visible absorbance spectra of CrinOCP2(ECH) and GcapOCP2(ECH) as compared with SynOCP1(ECH) (the difference spectra OCP1 minus OCP2 are also shown). The spectra were recorded from the chromatographic peak apex during spectrochromatography at 20 °C. The main maxima are indicated in nm. **E**. Superposition of CrinOCP2 and GcapOCP2 crystallographic monomers colored according to atomic B-factors from blue (low) to orange-red (high). Note the different position of the C-terminal tail (CTT) in the two structures, with a hinge Ala306 residue (magenta sphere). The domains (NTD, CTD), the N-terminal extension and the linker are shown.

The selected CrinOCP2 and GcapOCP2 amino acid sequences are 75% identical to each other (Fig. 1B), only ∼62 % identical to the model OCP1 from *Synechocystis* sp. PCC 6803 (SynOCP1), and even less, 52-54% identical to the recently characterized primordial OCP3 from *Gloeobacter kilaueensis* JS1 (GbacOCP3) [30]. The sequence identity relative to OCP2 from *Tolypothrix* sp. PCC 7601 (TolyOCP2) is 78% for CrinOCP2 and 84% for GcapOCP2, reflecting their different positions on the phylogenetic tree (Fig. 1A; full tree is presented in Supplementary Fig. 1). Curiously, the two selected OCP2 variants differed by the presence of the OCP2-specific Gln155 residue replacing the physiologically important Arg155 [11] found in OCP1 and OCP3 (vide infra).

### Heterologous OCP2 expression in ketocarotenoid-synthesizing *E.coli*

OCP2 proteins were reconstituted in *Escherichia coli* cells capable of ketocarotenoid synthesis [46–48] and contained no tags in the final form. For partially purified OCP2, analytical size-exclusion chromatography (SEC) revealed the main peak of monomers and minor dimeric and higher-order oligomeric peaks (Fig. 1C and Supplementary Fig. 2).

The oligomeric state of OCP remains controversial. First reports described OCP1° as a dimer [7] or a monomer [8,10,47]. Later reports suggested that the dimeric or monomeric state is an OCP family-dependent signature, with OCP1 and OCP3 being dimeric and OCP2 being monomeric [28,29]. SynOCP1 dimers were proposed to be stabilized by Arg27-Asp19 salt bridges [28,29]. Arg27 is lacking in CrinOCP2 (Ala), GcapOCP2 (Ser) and TolyOCP2 (Ser) but also in GbacOCP3 (it has Glu27 and Ser19). Nevertheless, our recent comparative SEC-MALS analysis of SynOCP1 and GbacOCP3 revealed that both proteins remain monomeric up to 700 μM [30], despite OCP1, for which most of the crystal structures were obtained (Table 1), so far crystallized exclusively as an antiparallel dimer stabilized by salt bridges involving Arg27.

In our hands, like previously for SynOCP1 and GbacOCP3 [30], analytical SEC revealed that both novel purified OCP2 holoproteins elute as single symmetrical peaks with unchanged positions even at very high protein concentration (up to 15.5 mg/ml (∼430 μM), see Fig. 1C and Supplementary Fig. 2). SEC-MALS analysis confirmed that both purified OCP2 holoproteins are monomeric and monodisperse (Fig. 1C and Supplementary Fig. 2), indicating that their higher-order oligomeric forms are very unstable.

HPLC analysis indicated that both OCP2 variants contained ECH as the predominant carotenoid (Supplementary Fig. 3). Absorbance spectra of these novel proteins drastically differed from SynOCP1 by a pronounced fine structure, reduced half-widths (e.g., 88 vs 105 nm for GcapOCP2 vs SynOCP1) and blue-shifted positions of the main peak maxima, the most extreme for GcapOCP2 (Fig. 1D). As a result, the difference “OCP1-minus-OCP2” spectra revealed the absence of a large part of absorbance in the red flange in OCP2 proteins, much more pronounced than “TolyOCP1-minus-TolyOCP2” [32]. Understanding of the underlying mechanisms clearly required structural analysis.

### Crystallization of two first OCP2 proteins

OCP2 crystallization proved difficult due to unknown reasons [32]. The amino acid substitution E25A improved GbacOCP3 crystallization [30], while mimicking alanine in this position found in OCP representatives in all families (in other *Gloeobacteria* OCP3a and SynOCP1 in particular [30]) and in reconstructed ancestral OCP sequences [41]. Since GbacOCP3 properties were insensitive to the E25A mutation [30], for OCP2 crystallization we also used this justified approach.

CrinOCP2 crystallized in many conditions, which led to structure determination at 2.2 A resolution (Table 2). The crystal structure revealed two CrinOCP2 molecules in the asymmetric unit (superimposable with the root mean square deviation (RMSD) of 0.44 A for 261 aligned Cα atoms (Supplementary Fig. 4)), but their relative position was different from the conserved antiparallel OCP1 dimer (Table 1 and Supplementary Fig. 3). The largest intermolecular interface 860.7 A^2^ (5.6% of solvent accessible surface area (SASA) of the monomer) featured few polar contacts, is only slightly larger than the largest intermolecular interface in GbacOCP3 structure (PDB ID: 8A0H, 768.6 A^2^, or 5.1% of the buried surface area [30]) and is much smaller than the common OCP1 interface (Supplementary Fig. 3 and Table 1). Hence, the crystallographic CrinOCP2 dimer is unlikely to be biologically relevant.

**Table 2.** Diffraction data collection and refinement statistics for GcapOCP2 and CrinOCP2.

| **Data collection** |  |  |
| --- | --- | --- |
|  | GcapOCP2 | CrinOCP2 |
| Diffraction source | BL17B1 beamline,<br>SSRF | Rigaku OD XtaLAB<br>Synergy-S |
| Wavelength (Å) | 0.98 | 1.54 |
| Crystal-to-detector distance (mm) | 230.0 | 42.6 |
| Rotation range per image (°) | 0.5 | 0.4 |
| Total rotation range (°) | 360 | 210 |
| Space group | P4<sub>3</sub>2<sub>1</sub>2 | P2<sub>1</sub>2<sub>1</sub>2<sub>1</sub> |
| a,b,c, (Å) | 52.66, 52.66, 216.83, | 60.05, 103.73, 109.62 |
| Resolution range (Å) | 54.21-1.80 (1.83-1.80) | 22.20–2.20 (2.27–<br>2.20) |
| Unique reflections | 29081 (1584) | 35514 (2978) |
| Completeness (%) | 98.2 (92.3) | 99.7 (98.2) |
| Average redundancy | 4.3 (4.4) | 7.5 (7.8) |
| I/sigma | 6.6 (0.7) | 12.3 (5.2) |
| CC<sub>1/2</sub> (%) | 99.6 (28.3) | 99.3 (87.4) |
| R<sub>pim</sub> (%) | 7.2 (141.4) | 5.1 (14.6) |
| **Refinement** |  |  |
| Reflections in refinement | 28909 | 35420 |
| R<sub>work</sub> (%) | 19.7 | 17.7 |
| R<sub>free</sub> (%) | 24.8 | 22.7 |
| RMS bonds (Å) | 0.01 | 0.01 |
| RMS angles (Å) | 1.82 | 2.05 |
| **Ramachandran plot** |  |  |
| Ramachandran favored (%) | 98.3 | 95.8 |
| Ramachandran allowed (%) | 1.7 | 3.4 |
| **Number of atoms** |  |  |
| Protein | 2375 | 4896 |
| Carotenoid | 41 | 82 |
| Other ligands | 29 | 4 |
| Solvent | 96 | 303 |
| **Average B-factors (Å<sup>2</sup>)** |  |  |
| Protein | 34.74 | 20.35 |
| Carotenoid | 27.35 | 12.31 |
| Other ligands | 53.35 | 37.63 |
| Solvent | 37.64 | 20.69 |
| MolProbity score | 1.48 | 1.88 |
| **PDB ID** |  |  |
|  | 8PZK | 8PYH |

Despite extensive crystal screening (>1000 conditions), GcapOCP2 formed just one small crystal, which fortunately was sufficient for structure determination using the synchrotron X-ray radiation. While the structure refined at 1.8 A revealed GcapOCP2 monomer in the asymmetric unit, two neighboring monomers in the crystal lattice formed a rather developed interface of 1072.8 A^2^ (7.3% of SASA, Supplementary Fig. 4), not superimposable with the crystallographic dimer of CrinOCP2 (RMSD >27 A for 595 Cα atoms). Nonetheless, both OCP2 were monomeric in solution up to at least 400 μM concentration (Fig. 1C and Supplementary Fig. 2).

### Comparative structural analysis of OCP1, OCP2 and OCP3

Determination of the first OCP2 structures completed the portfolio of structures for all three OCP families, enabling direct structural comparison of OCP1, OCP2 and OCP3. The overall architecture of the OCP2 monomer (Fig. 1E and Supplementary Fig. 4) is similar to SynOCP1 or GbacOCP3 (Ca RMSD is ≤1 A, Supplementary Fig. 5). The CTT of GcapOCP2 adopted an unusual conformation almost perpendicular to the long axis of the OCP molecule (Fig. 1E). Interestingly, while the CTTs of two neighboring GcapOCP2 molecules contacted each other, via hydrophobic interactions, the usual docking site for CTT on the CTD surface remained unoccupied. In contrast, in CrinOCP2 the CTT adopted the usual position on the CTD surface (Fig. 1E), as in other OCP crystal structures (Table 1). High B-factors of the CTT in GcapOCP2 structure implied its increased dynamics, coherent with the known role of this element in the carotenoid uptake and OCP photoactivation processes [49,50]. Comparison of the CTT position in GcapOCP2 and in SynOCP1 (or CrinOCP2) suggests that this element is rotated over Ala306 (Ala307 in SynOCP1/CrinOCP2) from its CTD-attached position (like in SynOCP1/CrinOCP2) to the outward position uniquely observed in GcapOCP2, and Glu311 (Glu310) appears to fix either position by polar contacts (Fig. 2A). The CTD surface for the CTT attachment is the same in GcapOCP2, CrinOCP2 or SynOCP1, hinting at the commonality of the identified CTT motion.

**Fig. 2.**
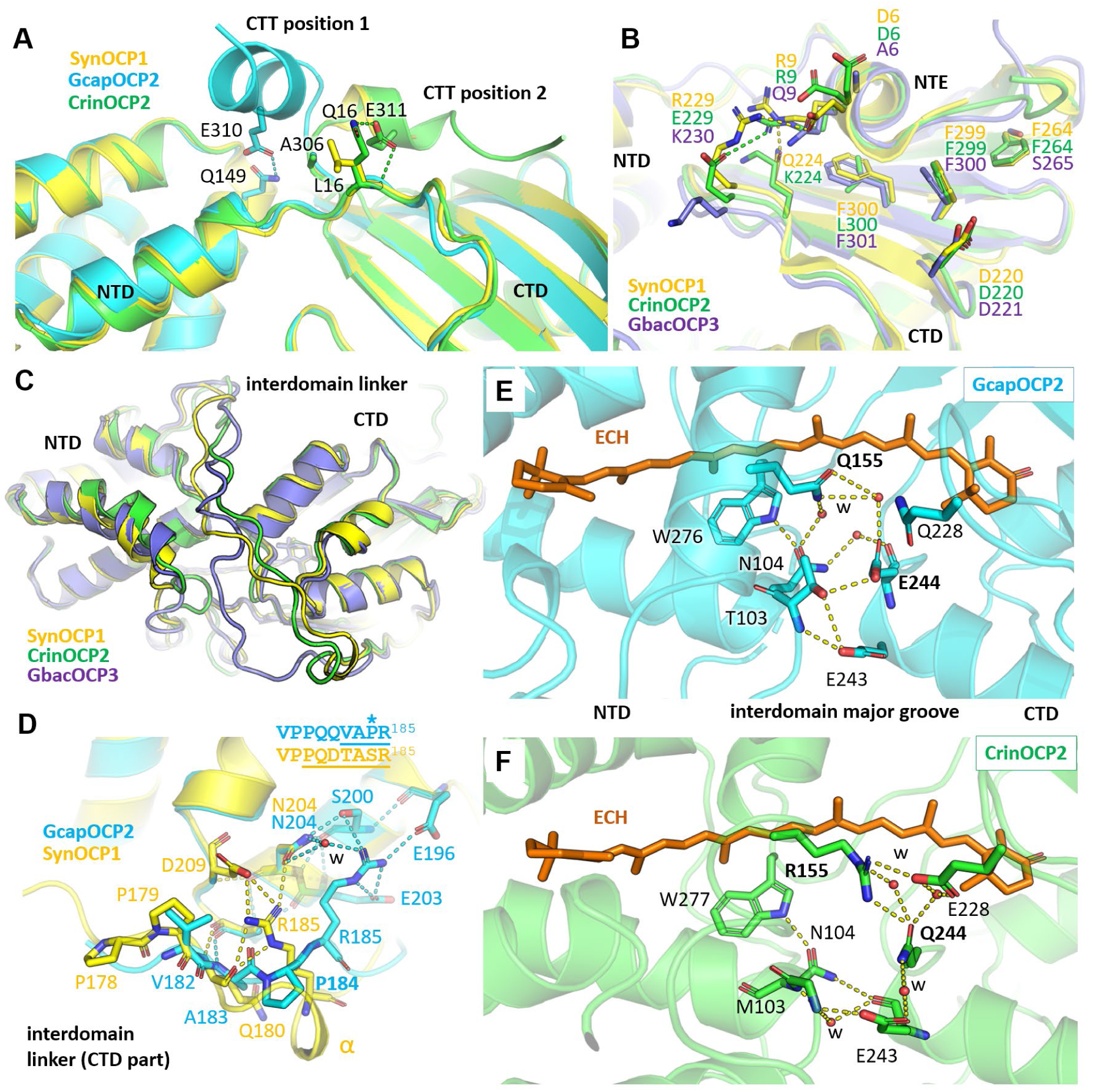
Structural features of OCP1, OCP2 and OCP3. **A**. Stabilization of the CTT in two different conformations inferred from the superposition of the crystal structures of SynOCP1, CrinOCP2 (CTT attached to the CTD) and GcapOCP2 (CTT sticks out). **B**. The conserved attachment of the NTE on the CTD of SynOCP1, CrinOCP2 and GbacOCP3. Polar contacts are shown by dashed lines colored according to the chain color. **C**. Superposition of the SynOCP1, CrinOCP2 and GbacOCP3 monomers showing the distinctly different conformations of the interdomain linkers. **D**. The altered linker conformation in GcapOCP2 dictated by its specific Pro184 causing a 3-residue positional shift back toward the interdomain groove. In the absence of Pro184, SynOCP1 (and CrinOCP2) features a 1-turn alpha-helix before Arg185. The shift results in Arg185 of GcapOCP2 making an alternative network of stabilizing interactions with the CTD residues. Polar contacts are shown by dashed lines colored according to the chain color. **E,F**. Specific interdomain interactions at the major groove dictated by the presence of either Gln or Arg in position 155 (NTD) and Gln or Glu in position 244 (CTD). Polar contacts are shown by yellow dashed lines. Waters (w) are shown by small red spheres.

NTE is connected similarly in OCP1, OCP2 and OCP3 (Fig. 2B). In CrinOCP2, Gln224 found in SynOCP1 and GcapOCP2, is replaced by Lys224, which is unable to connect with Arg9 of NTE found in all these three OCPs. In GbacOCP3, this polar contact is realized between Gln225 (CTD) and Gln9 (NTE). Another variation is the change of the conserved Phe300 residue found in SynOCP1 (Phe300), GcapOCP2 (Phe299) and GbacOCP3 (Phe301) to the non-aromatic Leu300 in CrinOCP2. Perhaps these variations weaken NTE attachment in CrinOCP2; but CrinOCP2 has Gln16 in place of Leu16 in the loop connecting the first α-helix of the NTE with the NTD, and this Gln16 provides extra stabilizing polar contact with Glu311 on the CTT (absent in SynOCP1 due to Leu16) (Fig. 2A). GcapOCP2 has the same Gln16-Glu311 combination (Glu310) and can likely form this bond, although it is not realized in GcapOCP2 structure because of the displaced CTT (Fig. 2A).

Despite the complete charge reversal due to the replacement of Arg229 of SynOCP1 by Glu229 of CrinOCP2 and GcapOCP2, it does not apparently affect the NTE attachment on the CTD because this sequence variation simply rearranges the salt bridges fixing the NTE on the CTD. As a result, the Arg229-Asp6 bridge in SynOCP1 is changed to Glu229-Arg9 bridge in both selected OCP2 variants. GbacOCP3 does not exhibit an equivalent salt bridge despite the presence of Lys229 and Gln9 combination (Fig. 2B), instead it features an intricate NTE/CTT locking mechanism stabilizing the dark-adapted conformation [30].

The interdomain linker is the most variable part in OCP (Fig. 1B) contributing to the functional signatures of distinct OCP types [29]. In GcapOCP2, the linker residues 166-179 could not be resolved due to the disorder, while at least in one CrinOCP2 monomer (chain B) the whole linker could be traced in the electron density. This allowed us to appreciate the different conformations of the full linkers in OCP1, OCP2 and OCP3 (Fig. 2C). The traced part of the GcapOCP2 linker reveals the peculiar difference around Pro184, which is absent in SynOCP1 and CrinOCP2 and is replaced by a short helical element, in turn absent in the GcapOCP2 structure (Fig. 2D). This Pro, present also in TolyOCP2, shifts the conserved EPXVPP linker segment of GcapOCP2 three residues toward the major interdomain groove region (Fig. 2D), forcing Arg185 of GcapOCP2 to form an alternative H-bond network compared to the equivalent Arg185 residue of SynOCP1 (or CrinOCP2). Nevertheless, the linker-immobilizing role of this arginine is preserved in all three cases, as supported by the compromised ability of R185A/E mutants to undergo R-O transition [29]. Although the consequences of the Pro184-associated register shift observed for GcapOCP2 is not fully clear, such details are hardly predictable by Alphafold [51], which yields a structural model with as large as ∼2 A Cα RMSD from with the refined GcapOCP2 structure. This underlines the importance of obtaining experimental structures of OCP.

Interdomain contacts in the major OCP2 groove are noteworthy (Fig. 2E and F). The canonical SynOCP1 has the salt bridge between Arg155 in the NTD and Glu244 in the CTD, stabilizing the domain interface and breaking up during OCP photoactivation and PBS quenching [8,11]. GbacOCP3 has this salt bridge too [30]. Yet, Arg155 and Glu244 are not strictly conserved across the OCP families: Arg155 is replaced by Gln155 in most of OCP2 members including GcapOCP2, which puzzled researchers due to the known role of Arg155 [28]. In the GcapOCP2 structure, Q155 does not form salt bridges – only water-mediated contacts with the CTD (Fig. 2E). The only direct H-bonds in the GcapOCP2 groove involve Thr103, which connects to Glu243 and Glu244 simultaneously. Both SynOCP1 and GcapOCP2 contain Glu244, but only SynOCP1 has the typical Arg155-Glu244 salt bridge [8]. While CrinOCP2 contains Gln244, it also salt bridges with Arg155. In addition, Arg155 of CrinOCP2 makes direct connection with Glu228 of the CTD, further tightening the domain interface, while Gln228 in SynOCP1 and GcapOCP2 cannot reach the counterpart Arg155 (SynOCP1) or Gln155 (GcapOCP2), which are >4.5 and >5.5 A away, respectively (Fig. 2E,F). Therefore, the interdomain interface in CrinOCP2 is tighter than in GcapOCP2, which harbors the OCP2-specific Gln155.

### Protein-carotenoid interactions dictate spectral signatures of OCP1, OCP2 and OCP3

Despite the apparent similarity of the conformation and molecular environment of the ECH molecule in OCP2 and in OCP1, several differences are notable (Fig. 3A): i) two alternative conformations of Trp287 in GcapOCP2, ii) the presence of Met205 in both CrinOCP2 and GcapOCP2 instead of leucine in SynOCP1, iii) a different conformation of Phe278 in CrinOCP2 (Ala277 in GcapOCP2), iv) the presence of polar Ser114 in CrinOCP2 and GcapOCP2 instead of Gly in SynOCP1 and v) the OCP2-specific Gln155 instead of Arg155 in SynOCP1 (and CrinOCP2). Careful inspection of residues directly interacting with the keto group of ECH in various OCP structures (Fig. 3B) revealed the determinants of the spectral features (Fig. 3C,D). While in crystals of SynOCP1 the keto group of ECH forms two H-bonds with Tyr201 (2.6 A) and Trp288 (2.9 A) and these contacts are also present in OCP2 and OCP3, the corresponding distances are slightly different in GcapOCP2 (Fig. 3B) and CrinOCP2 (Supplementary Fig. 6). In GcapOCP2, the H-bond with Tyr201 is subtly shorter (2.5 A) and with Trp287 is subtly longer (3.0 A) than in SynOCP1, and we also observed an alternative Trp287 conformation. This is reminiscent of the two Trp201 conformations found in the SynOCP1^WW^ mutant structure (PDB ID: 6T6K) (Fig. 3B), in which the distance to the nitrogen atom of Trp201 also increased (to 3.2A) [19]. Strikingly, the visible absorbance spectra of GcapOCP2(ECH) and SynOCP1^WW^(ECH) virtually coincide and have the pronounced fine structure (Fig. 3C) indicative of strong coupling of vibrational and electronic states [34]. We assume that effectively one H-bond donor, Trp288 in SynOCP1^WW^(ECH) or Tyr201 in GcapOCP2(ECH), yields the pronounced fine structure of the absorbance spectrum (Fig. 3C), as the same spectral shape is displayed by the SynOCP1^W288A^ mutant with Tyr201 as the sole H-bond donor in the CTD [19] (Fig. 3C) or by the SynOCP1^Y201F^ mutant with the single H-bond to Trp288 [21].

**Fig. 3.**
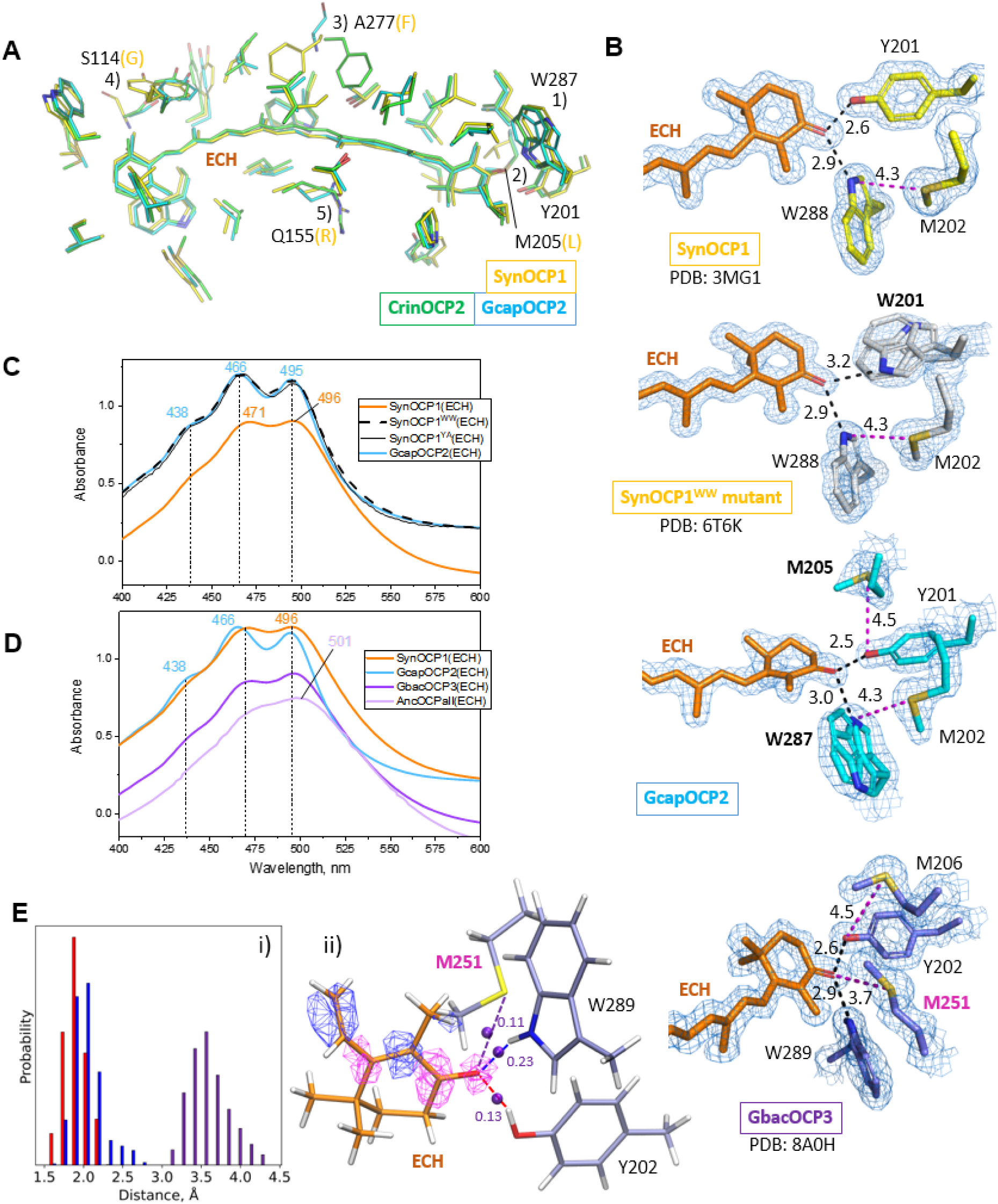
OCP-carotenoid interactions determine the spectral properties. **A**. Superposition of the carotenoid-binding sites of SynOCP1, CrinOCP2 and GcapOCP2 reveals the number of features (from 1 to 5: OCP2-specific residues (black font) are compared with the residues of SynOCP1 (yellow font)). Residues having atoms within 4 A of the echinenone atoms are shown by thin sticks following the protein color coding. Echinenone (ECH) is shown by orange sticks. **B**. Residues directly contacting the carotenoid in the CTD of SynOCP1 WT, SynOCP1 Y201W mutant (WW), GcapOCP2 or GbacOCP3. 2Fo-Fc electron density maps contoured at 1.5σ are shown. H-bonds with the carotenoid keto group are shown by black dashed lines, potential chalcogen bonds involving the adjacent Met residues are shown by magenta dashed lines. The bond lengths are shown in angstrom. **C**. Normalized absorbance spectra of SynOCP1(ECH) or its mutants Y201A (WW) and W288A (YA) with GcapOCP2(ECH). **D**. Normalized absorbance spectra of SynOCP1(ECH), GcapOCP2(ECH), GbacOCP3(ECH) and the spectrum of the reconstructed ancestor of all OCPs (AncOCPall; digitized from [41]). The spectra are shifted along the Y axis for convenience. The main peak maxima are shown in nm. **E**. QM/MM model of GbacOCP3. i) distributions of distances between the ECH keto oxygen and hydrogen atoms of Y202 (red bars) and W289 (blue bars); and sulfur atom of M251 (lavender bars) along the QM/MM MD trajectory. ii) The structure at the minimum on the potential energy surface showing interatomic interactions of the keto oxygen and other atoms (dashed lines) with the corresponding electron density values (in a.u.) at bond critical points (violet balls). Magenta and blue isosurfaces (+/-0.005 a.u.) depict calculated electron density increase and decrease, respectively, upon electronic excitation.

The absorbance spectrum of the wild-type SynOCP1(ECH) is slightly red-shifted and broadened (Fig. 3C), which is likely due to the two effective H-bond donors contacting the ECH keto group (Fig. 3B). Following this logic, we assumed that further broadening and red-shift, such as displayed by OCP3 (Fig. 3D), reflects an increased number of possible bonds involving the carotenoid keto group. Scrupulous inspection of our previously solved GbacOCP3 structure (PDB ID: 8A0H [30]) revealed a unique Met251 residue in vicinity of the ECH keto group (not found in OCP1 or OCP2), bringing the ECH oxygen atom and the sulfur atom of M251 to a ∼3.7 A distance (Fig. 3B). Tyr202, Trp289 and Met251 triangulate the position of the ECH keto oxygen, hinting at the direct bonding interaction with Met251 (Fig. 3B). We performed molecular dynamics simulations with combined quantum mechanics/molecular mechanics (QM/MM) potentials and the following energy minimization to analyze the ECH keto group oxygen interactions (Fig. 3E, i and ii). The QTAIM analysis demonstrated the presence of the interactions between the ECH oxygen and the sulfur atom of the Met251 side chain, as there was a bond critical point between these two atoms. To quantify these interactions, we calculated electron density values corresponding to hydrogen bonds with Tyr201 and Trp289 and a chalcogen bond with the Met251 sulfur atom. Importantly, all these bond critical points had comparable values of electron densities (Fig. 3E, ii) that explained the significant contribution of the chalcogen bond. Distribution of this bond is centered at around ∼3.6 A (Fig. 3E, i), which is close to the corresponding distance in GbacOCP3 (PDB ID: 8A0H). The electron density redistribution upon electronic excitation was calculated to explain the importance of keto oxygen interactions with H-bond donors/σ-hole containing atoms (Fig. 3E, ii). Upon excitation, the electron density increases on the keto oxygen or, in other words, the charge becomes more negative on this atom. Therefore, interactions mentioned above stabilize the excited state that is manifested in the experimentally observed red-shift of the absorption spectrum.

The discovered chalcogen bond explains the spectral signatures of OCP3 members more generally. In fact, Met dominates in the position 251 in the whole OCP3 family and is also present in the reconstructed ancestor of all OCP proteins (AncOCPall), which has even a slightly more red-shifted and broadened absorbance spectrum [41] (Fig. 3D). Furthermore, the recently reported crystal structure (PDB ID: 7YTF) of OCPX2 from *N. flagelliforme* (OCP3b, according to the recently proposed classification [30], see Supplementary Fig. 1) also reveals the same triangulation of the carotenoid keto group by Tyr202, Trp289 and Met251. A shorter chalcogen bond (3.5 A vs 3.7 A in GbacOCP3 structure) nicely correlates with a more red-shifted and broadened absorbance spectrum of *N. flagelliforme* OCP3b [31], which is very similar to that of AncOCPall [41] (Fig. 3D). While many other proteins, including photoreceptors, have methionines at a sensible distance for making chalcogen bonds with oxygen atoms of their ligands, these bond types remain largely overlooked, even though they can be physiologically relevant (Supplementary Table 1).

### Absorbance spectrum and photocycle signatures of OCP1, OCP2 and OCP3

In solution, the dark-adapted SynOCP1 contains a significant amount of the spectrally red state (state 1) formed due to spontaneous H-bonds breaking [19]. During relaxation of the red form, SynOCP1 first forms an “orange” state intermediate (state 2) lacking the fine structure of the carotenoid absorption spectrum and featuring ketocarotenoid interaction with both H-bond donors [52]. Further relaxation leads to an increased role of the ketocarotenoid interaction with only one H-bond donor (Y201 or W288), which leads to the manifestation of the fine absorption structure (state 3) [52]. Thus, the absorption of the compact dark-adapted state of various OCP members (Fig. 4A-D) can be represented by the sum of states (1, 2 and 3) reflecting different interaction patterns of the carotenoid and the protein environment. Such decomposition enables assignment of these states using the available structural information (see Fig. 3B).

**Fig. 4.**
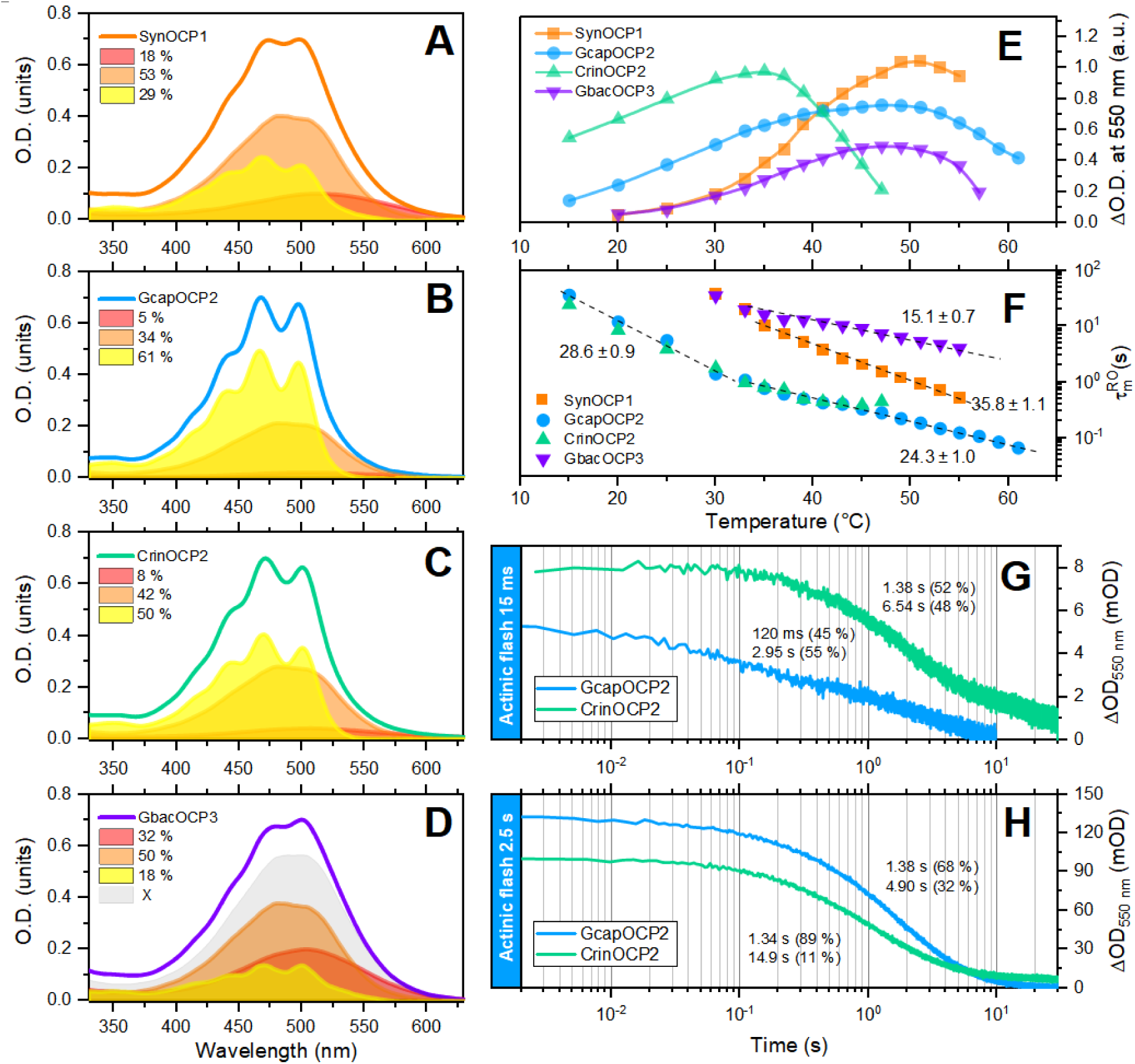
Spectral characteristics and photocyclic transformations of OCP1, OCP2 and OCP3 members. Absorption spectra and results of component analysis of spectra of SynOCP1 (**A**), GcapOCP2 (**B**), CrinOCP2 (**C**) and GbacOCP3 (**D**) preparations adapted to darkness at 5 °C. X represents the sum of non-vibronic components in the GbacOCP3 spectrum. (**E**) – Relative yields of OCP photoactivation photoproducts in response to a 100 ms actinic flash (445 nm) presented as a function of temperature. Yields were measured as changes in OD_550_. Protein concentrations were selected so that the absorbance of all samples ranged from 0.43 to 0.48 OD units at 445 nm. (**F**) – temperature dependence of the lifetime of the dominant kinetic component of the relaxation of red forms after 100 ms actinic flash. Numbers correspond to estimates of the activation energy of the transition in kcal/mol. Relaxation kinetics of OCP red forms measured as changes in optical density at 550 nm in response to an actinic flash of 15 ms (**G**) and 2500 ms (**H**). Time zero corresponds to the moment when the actinic flash is turned off. Experiments were performed at 30 °C. The concentration of protein preparations was equalized by the carotenoid absorbance in the visible part of the spectrum and was ∼5 μm. Numbers indicate the characteristic durations of the kinetic components and their contributions, estimated by approximating the experimental data by the sum of two decaying exponentials.

Although being the narrowest compared to OCP1 and OCP3, the absorption spectra of dark-adapted OCP2 are also broadened due to the presence of several carotenoid states (Fig. 4A-D). In CrinOCP2 (Fig. 4B) and especially GcapOCP2 (Fig. 4C), the contribution of the state (3) with a pronounced vibronic structure and a single H-bond is the predominant one. This is in line with the GcapOCP2 structure, in which Trp287 is in the alternative conformation unable to interact with the carotenoid, increasing the probability of interaction with Y201. By contrast, GbacOCP3, characterized not only by developed protein-protein contacts stabilizing the compact state [30] but also by the additional carotenoid contact with Met251, has the least fraction of the vibronic state (Fig. 4D). Given the presence of three potential bonds (with Y, W and M), the decomposition of the dark-adapted OCP3 spectrum into states 1-3 like for OCP1 and OCP2 is a simplification, since we do not yet know the spectra of OCP3-specific states in which the carotenoid interacts only with methionine or the methionine-tyrosine/methionine-tryptophan pair. Nevertheless, the total fraction of the states devoid of any fine structure is largest in GbacOCP3 (“X” in Fig. 4D), which is likely associated with the increased heterogeneity imposed by the Met251 presence.

Both novel OCP2 species were photoactivatable, however, actinic flashes even at high power and low temperature were not able to completely convert them to the red form, which hinted at the high relaxation rate of the red forms (Supplementary Fig. 7). This unusual behavior was investigated further at different temperatures (Fig. 4E and F).

We accidentally found that CrinOCP2 absorbance spectrum changes with increasing temperature even in the dark (Supplementary Fig. 8), resembling the chemical SynOCP1 activation by high concentrations of NaCSN [53], and it was not reversible within 1 h after cooling. Nevertheless, the absorption spectra of the protein even at 52 °C had very low light scattering, suggesting protein aggregation was not the case. Moreover, such a spectrally red CrinOCP2 form decreased fluorescence intensity of *Synechocystis* PBS by ∼17% and lifetime by ∼5%, although this was much less than high-light-induced quenching (vide infra).

Since the OCP1 photocycle conceals a variety of intermediates [19,20], with the most relevant for the regulation of non-photochemical quenching occurring on the ms scale, we conducted similar studies for OCP2. At short flashes (15 ms), the relaxation kinetics of the red forms of CrinOCP2 and GcapOCP2 differed significantly. At 30 °C in the relaxation kinetics of photoactivated GcapOCP2, a significant contribution belongs to components with lifetimes of <120 ms, but under identical conditions the CrinOCP2’s main component lifetime is ∼10 times longer (Fig. 4G). With increasing actinic flash duration, the component composition of the relaxation kinetics of photoactivated GcapOCP2 changes so that fast components are replaced by slower ones (Fig. 4H). This behavior is generally consistent with accepted models of OCP1. However, in the case of CrinOCP2, the main component even with long flashes remains 1.34 s and, therefore, likely represents the final red form (Fig. 4H). Thus, in CrinOCP2 the final red state is formed much earlier than in GcapOCP2, and its photocycle probably bypasses some intermediates [20,52]. Indeed, only at 20 °C we detected the fast component (120 ms) in the relaxation kinetics of photoactivated CrinOCP2 after 100 ms actinic flash (Supplementary Fig. 9).

Amplitudes of the OD_550_ change in solutions of different OCPs with the same concentration in response to identical actinic flashes allowed us to compare the quantum yield of photoproduct formation (Fig. 4E). Importantly, both OCP2 representatives had higher photoproduct yields than GbacOCP3 in a wide temperature range, indicating that the abundance of states with a vibronic structure of the absorption spectrum is crucial for the photoactivation of OCPs. When considering these dependencies, it should be taken into account that the observed amplitude of OD_550_ changes generally depends on the lifetime of red states and the temporal resolution of the registration system. Thus, when using 100 ms flashes, we inevitably lose information about shorter-lived intermediates (with a potentially much larger yield) and focus on the final (longer) stages of the photocycle. We would like to note that taking into account the small contribution of the photosensitive state compared to OCP2, the quantum yield of SynOCP1 is high, which is probably due to the significantly lower activation energy for its O-R transition (Table 3). Together with the long lifetime of the red forms this makes SynOCP1 an almost irreversible and thereby poorly controlled regulator of photoprotective reactions in the absence of FRP.

**Table 3.** Comparative characteristics of representatives of the three OCP families.

| Property | SynOCP1 | GcapOCP2 | CrinOCP2 | GbacOCP3 |
| --- | --- | --- | --- | --- |
| Initial photoactivatable fraction (with vibronic structure) | 29 % | 61 % | 50 % | 18 % |
| Red fraction in dark-adapted sample | 10-25 % | 6 % | 8 % | 32 % |
| Temperature with maximum yield of photoproducts, °C | 52 | 48 | 35 | 47 |
| Yield of photoproducts | High | High | High | Very low |
| $E_a^{O-R}$, kcal/mol | $7.5 \pm 0.3$ [19] | $14.8 \pm 0.5$ | $19.2 \pm 1.9$ | $16.1 \pm 0.6$ |
| Intermediates after 100 ms actinic flash at 30 °C | Yes | Yes | No | No |
| Red states relaxation at 5 °C | Very slow | Fast | Fast | Fast |
| $E_a^{R-O}$, kcal/mol | $35.8 \pm 1.1$ | $24.3 \pm 1.0$ | $28.7 \pm 0.9$ | $15.8 \pm 0.7$<br>($21.8 \pm 1.9$) |
| Interaction with SynFRP | Tight binding,<br>great acceleration<br>of R-O | Weak binding<br>Deceleration | Weak binding<br>Deceleration | No effect |
| PBS quenching | Yes | Yes (relatively<br>inefficient) | Yes | Yes |

Temperature dependencies of OCP2 kinetics reveal fundamental differences in the functioning of CrinOCP2 and GcapOCP2 (Fig. 4E): the optimum temperature for photoactivation of GcapOCP2 is comparable to that of SynOCP1 (48-50 °C), while that of CrinOCP2 is, conversely, much lower (35 °C). This is probably a reflection of the optimal physiological conditions for the respective cyanobacterial species – the optimal growth temperature of *Crinalium* is ∼25 °C [43], while *Gloeocapsa* PCC 7428 inhabits moderate hot springs with an apparently higher temperature [45]. Therefore, we attribute the appearance of long-lived components of relaxation kinetics (Fig. 4G, H) in all OCPs at elevated temperatures (especially in CrinOCP2) to protein misfolding and interaction of the red form dimers. Comparison of the average relaxation rates of red forms in OCP1 and OCP2 shows that GcapOCP2 relaxes much faster than OCP1, which potentially makes it a convenient target for studying the initial stages of the photocycle at elevated temperatures (up to 45 °C). At the same time, CrinOCP2 relaxes significantly faster than SynOCP1, GbacOCP3 and GcapOCP2 at reduced temperatures. As already noted, the absence of FRP genes in OCP2 family members requires alternative ways of regulating non-photochemical quenching. Perhaps, one way is to reduce the probability of photoactivation like in GbacOCP3, while another way to accelerate adaptation to changing conditions is to accelerate photocyclic transitions like in OCP2. However, this does not exclude the potential possibility of OCP2 interaction with FRP.

### Interaction of OCP1, OCP2, OCP3 with FRP and the phycobilisome

The R-O transition of various OCPs was studied upon photoactivating the proteins at different temperatures by continuous blue LED illumination until reaching the equilibrium, then switching off the LED and following OD_550_ changes with or without SynFRP. τ_1/2_ for the R-O transition of GcapOCP2 alone decreased by two orders of magnitude from 203 s (5°C) to 8 s (23°C), whereas the presence of a two-fold molar excess of SynFRP slightly increased τ_1/2_ at every temperature tested (e.g., from 203 to 250 s at 5°C and from 12 to 16 s at 20°C) (Supplementary Fig. 10), suggestive of the altered SynFRP interaction with GcapOCP2 decelerating its R-O transition. To our knowledge, the decelerating effect of FRP has not been addressed until very recently, with respect to SynFRP interaction with ancestral OCP [41]. Under the continuous photoactivation regime, for GcapOCP2, the *E*a^R-O^ of ∼26.5 kcal/mol inferred from the Arrhenius plot did not change in the presence of SynFRP, however, the line shifted parallel to that in the SynFRP absence (Fig. 5A), implying that with SynFRP, GcapOCP2 requires more attempts to join the domains and restore the OCP^O^ state [20]. The energy barrier (*E*a^R-O^ 26.5 kcal/mol) is similar to the one reported for GbacOCP3 (*E*a^R-O^ 25.7 kcal/mol [30]) and is much lower than that of SynOCP1 (*E*a^R-O^ 35.8 kcal/mol) (Table 3). Under the continuous photoactivation regime, the R-O rate of GcapOCP2 was slightly faster than that of GbacOCP3 and drastically faster than that of SynOCP1 at every temperature tested (Fig. 5B and Supplementary Fig. 10). The R-O conversion of CrinOCP2 was even faster than GcapOCP2 at temperatures 5-14°C (τ_1/2_: 140 vs 203 s at 5°C, 73 vs 103 s at 8°C, 40 vs 53 s at 11°C, 24 vs 31 s at 14°C), although we did not build Arrhenius plot for CrinOCP2 from these data due to the presence of dominating slow processes at temperatures above 17°C (Supplementary Fig. 10). The decelerating effect of SynFRP was noticeable, albeit smaller, also for CrinOCP2 (τ_1/2_ increased from 140 to 155 s at 5°C, from 73 to 83 s at 8°C, from 45 to 49 s at 11°C, from 24 to 28 s at 14°C; Supplementary Fig. 10), again suggestive of the counterproductive SynFRP/OCP2 interaction.

**Fig. 5.**
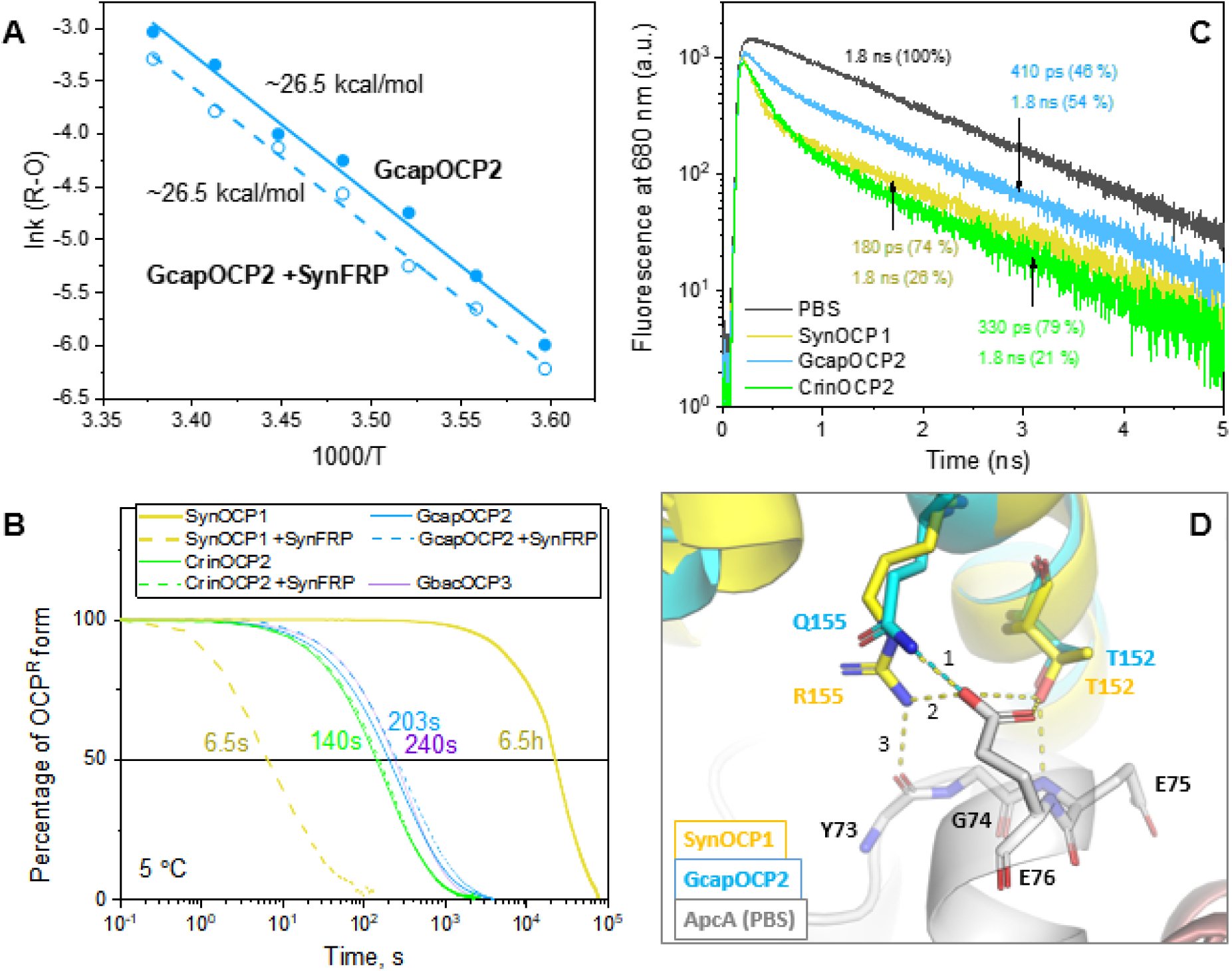
Functional properties of OCP2. **A**. Arrhenius plot showing the decelerating effect of SynFRP on the R-O transition of GcapOCP2 recorded after switching off continuous illumination (see Supplementary Fig. 10 for all kinetics). *E*a^R-O^ values for the absence and the presence of SynFRP are indicated. **B**. Normalized semi-log plot facilitating assessment of the R-O transition rates for OCP1, OCP2 and OCP3 and the effects of SynFRP. Time required for half R-O transition (τ_1/2_) is indicated for SynOCP1 (+/-SynFRP) and OCP2 and OCP3 without SynFRP, with the consistent color coding. Concentration of OCP was 10 μM, FRP – 20 μM. **C**. PBS fluorescence decay kinetics in the absence (black line) or in the presence of SynOCP1 (yellow line), CrinOCP2 (green line) or GcapOCP2 (blue line) in 0.75 M phosphate pH 7.0 containing 0.75 M sucrose at 5 °C. The ratio of OCP to PBS was ∼100. **D**. Superposition of the GcapOCP2 crystal structure on the NTD of SynOCP1 complexed with the *Synechocystis* phycobilisome (PDB ID: 7SC9 [27]) showing the difference in H-bonding network formed by Arg155 in SynOCP1 (present also in CrinOCP2) vs Gln155 in GcapOCP2. H-bonds are shown by dashed lines, yellow for Arg155 environment, cyan – for GcapOCP2. The effective H-bonds formed by Arg155 are numbered (only one H-bond is possible for Gln155).

By using normalization of the kinetic curves at a particular temperature (5°C) and their semi-log representation we could conveniently compare the rate of R-O transition for OCP1, OCP2 and OCP3, and the effect of SynFRP (Fig. 5B). While SynOCP1 is the slowest in terms of R-O relaxation (Fig. 5B and Supplementary Fig. 10), followed by GbacOCP3, GcapOCP2 and CrinOCP2, it becomes the fastest in the presence of SynFRP (τ_1/2_ drops grossly from 6.5 h to 6.5 s at 5°C). The latter either does not affect (GbacOCP3 [30]), or slightly decelerates the R-O transition (GbacOCP2 and CrinOCP2). Although looking close on the log scale, the kinetics of CrinOCP2 and GcapOCP2 are still appreciably different, hinting at the weaker interdomain interactions due to Gln155 in GcapOCP2 (Fig. 2E, F).

Since the residue-155 is important also for the interaction with PBS [11], we compared the PBS quenching ability of CrinOCP2, GcapOCP2 and SynOCP1 using the model *Synechocystis* phycobilisomes (SynPBS), whose complex with SynOCP1 has recently been solved [27]. Fluorescence decay kinetics of unquenched PBS are characterized by the characteristic fluorescence lifetime of 1.8 ns (Fig. 5C), whereas the photoactivation of any OCP tested resulted in significant PBS fluorescence quenching, as judged from the appearance of a fast component of the decay attributable to an OCP-quenched PBS state. Although the fast (∼200 ps) component of the kinetics itself does not characterize the rate of energy transfer between PBS and carotenoid in OCP, but rather the energy transfer between PBS rods and the core, we observed significant differences for OCPs tested. For SynOCP1(ECH) or CrinOCP2(ECH), as much as 75% of the initial PBS fluorescence was quenched, indicating the highly efficient binding of OCP into the PBS core. In contrast, in the case of GcapOCP2(ECH) only 50% of PBS fluorescence was quenched, suggesting its reduced binding efficiency to the PBS core. Upon inspection of the interface between SynOCP1 and SynPBS (PDB ID: 7SC9) and considering the multiple sequence alignment (Fig. 1B) we noticed that the main difference is around OCP2-specific Gln155 replacing the Arg155 of other OCPs (including CrinOCP2). Superposing the GcapOCP2 structure obtained in this work onto SynOCP1 in the SynOCP1-SynPBS structure (PDB ID: 7SC9), one can see that the maximum of three effective H-bonds that could be formed by Arg155 with the counterpart Glu76 side chain and the backbone oxygen of Tyr73 of the ApcA of PBS is reduced to only one in the case of shorter Gln155 in GcapOCP2 (Fig. 5D). This implies that the Arg155Gln replacement does not abolish the OCP2-PBS interaction but decreases its efficiency, in accordance with what is observed in the experiment (Fig. 5C) and with the data on a similar effect of the Arg155Lys substitution in SynOCP1 [11].

The observed deceleration of OCP2 R-O relaxation in the presence of SynFRP (Fig. 5A, B and Supplementary Fig. 10) may be due to the direct protein-protein interactions. Previous works showed that SynFRP forms tight complexes with SynOCP1 devoid of the NTE covering the FRP-binding site on the CTD surface [13,24]. The hybrid low-resolution structure of this complex (SASBDB ID: SASDDG9 [24]) is now validated by Alphafold, which yields the same models regardless of the NTE presence in SynOCP1 since FRP attachment to the CTD displaces the NTE from its binding site (Fig. 6A). Recruited by such binding on the CTD, SynFRP dimer is also known to interact with SynOCP1 forms with separated NTD and CTD, i.e. the W288A mutant mimicking the photoactivated OCP state [10], or the SynOCP1 apoform [54]. With the wild-type holoprotein, this was proposed to bring the domains together due to the complementarity of the electrostatic surfaces of OCP and FRP [24]. Interestingly, all Alphafold predictions of the SynOCP1/SynFRP complex were identical (Fig. 6A), whereas structural models of the GcapOCP2/SynFRP complex differed substantially by the conformation of GcapOCP2, which in some cases had separated NTD and CTD. Yet, the predicted position of the SynFRP head on the CTD remained similar, indicating that the main FRP-binding site is likely preserved (Fig. 6B).

**Fig. 6.**
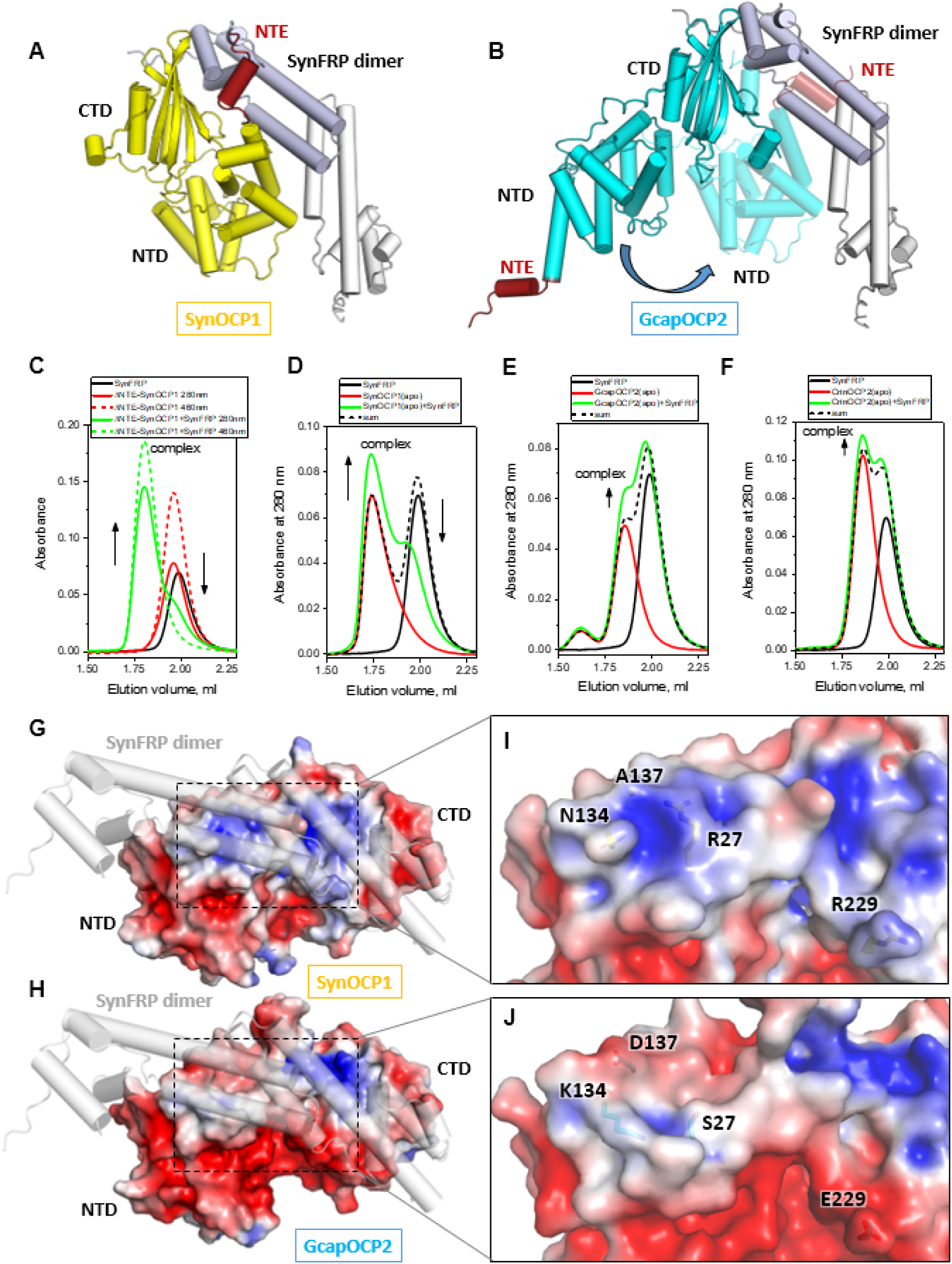
Interaction of OCP1 and OCP2 with SynFRP. **A**. An Alphafold model of the SynOCP1 complex with SynFRP dimer consistent with the low resolution hybrid structure (SASBDB ID: SASDDG9 [24]). Only one model is shown, others were identical. SynFRP binds via its head domain on the CTD displacing the NTE (red). **B**. Two Alphafold models of the GcapOCP2 complex with SynFRP representing GcapOCP2 with either separated or joined NTD and CTD. Imaginary rotation of the NTD over the linker from the extended to compact conformation is shown by the arrow. GcapOCP2 is shown by cyan ribbon, NTE is red. C-F. Interaction of SynFRP with ΔNTE-SynOCP1(ECH) (**C**), SynOCP1 WT apoform (**D**) or the apoforms of GcapOCP2 (**E**) and CrinOCP2 (**F**) studied by analytical SEC. Arrows illustrate complex formation. Black dashed lines on **D-F** represent algebraic sums of the SynFRP and OCP profiles (i.e., scenario with no interaction). **G,H**. The electrostatic potential distribution on the surface of the crystal structures of SynOCP1 (PDB ID: 3MG1) (panel **G**) and GcapOCP2 (this work) (panel **H**) calculated by APBS tools (color gradient from red (–3 kT/e) to blue (+3 kT/e)) at 150 mM salt concentration (i.e., matching the experiment conditions). SynOCP1 and GcapOCP2 monomers are overlaid onto the SynOCP1 chain of the Alphafold model of the SynOCP1/SynFRP dimer complex, to illustrate the difference of the electrostatic potentials within the FRP footprint (dashed rectangles). **I,J**. Close-up views of the FRP docking site on the NTD showing the location of the oppositely charged residues specific to OCP1 or OCP2.

Such interactions were directly tested by SEC (Fig. 6C-F), which first confirmed the formation of tight complexes between SynFRP and ΔNTE-SynOCP1(ECH) (Fig. 6C) or the apoform of SynOCP1 (Fig. 6D). Surprisingly, we also detected SynFRP interaction with the GcapOCP2 apoform (Fig. 6E) and, barely detectable, with the CrinOCP2 apoform (Fig. 6F). Intriguingly, the efficiency of SynFRP binding to OCP2 with separated domains somehow correlated with the degree of the decelerating effect of SynFRP on the OCP2’s R-O transition (GcapOCP2>CrinOCP2) (Fig. 5). Remarkably, in the center of the rather conserved CTD surface, only CrinOCP2 features Leu300 instead of Phe300 found in SynOCP1 or GcapOCP2, which can at least partly be responsible for the weaker SynFRP/CrinOCP2 interaction observed (Fig. 2B).

FRP was reported to bind to the resurrected ancestor of all OCPs (AncOCPall), decelerating its R-O transition [41]. This agrees with our results showing that SynFRP interacts with the expanded OCP2 forms but decelerates the R-O transition of OCP2, in contrast to its renowned accelerating effect on OCP1 (Fig. 6F). Therefore, while SynFRP can be recruited to the CTD of OCP2s, it cannot properly bridge the CTD and NTD together to accelerate R-O conversion and instead decelerates it to a different extent, depending on the protein tested. This implies that the secondary FRP binding site(s) on the OCP2 NTD are altered in the case of OCP2. Indeed, comparison of the electrostatic potential distribution on the surface of SynOCP1 or GcapOCP2 revealed striking differences within the SynFRP binding footprint on the NTD (Fig. 6G-J). The charge complementarity of the SynOCP1 and SynFRP surfaces at the evolutionarily cold region [24], including the positively charged area centered at Arg27 (Fig. 6G and I), is no longer maintained in GcapOCP2 due to the R27S and A137D amino acid changes, which collectively reverse the charge of the area from positive to negative (Fig. 6H and J). Even stronger charge reversal is found in CrinOCP2, where the same area is occupied not only by Ala27 and Asp137, but also by two additional negatively charged residues, Asp134 and Glu133 (Supplementary Fig. 11). Such disturbance of charge complementarity likely results in SynFRP preventing the OCP2 domain reassociation, which is reflected in deceleration of the R-O transition (Fig. 5). Notably, Arg27 and Ala137 (sometimes, Ser137) are hallmarks of the OCP1 clade, whereas OCP2 members almost uniformly contain Ala/Ser27 and Asp/Glu137 [41], which nicely correlates with the accelerating/decelerating effect of SynFRP on OCP1/OCP2. Furthermore, this logic is confirmed by the decelerating effect of SynFRP on AncOCPall, because this OCP variant has Arg27Leu (similar to OCP2), Ala137 (like OCP1) but has Asp134 which all the same reverses the charge of the FRP binding footprint on the NTD (Supplementary Fig. 11). This cause-and-effect relationship is additionally supported by the observation that GbacOCP3 is not regulated by and does not interact with SynFRP [30], because this protein correspondingly has i) Glu27 instead of Arg27 and Asp137 instead of Ala (like OCP2s tested), which reverses the charge on the NTD repelling SynFRP, and ii) contains a compromised SynFRP binding site on the CTD due to the presence of Ser265 instead of an aromatic Phe/Tyr found in OCP1, OCP2 (Fig. 2B) and AncOCPall (Fig. 1B), all of which are known to interact with SynFRP by their CTD [41]. The isolated CTD domain of GbacOCP3, neither in the apo nor in the carotenoid-bound form, cannot interact with FRP from two different cyanobacteria, *Synechocystis* and *Arthrospira*, providing explanation for the inability of FRP to dock onto the CTD surface of GbacOCP3 containing Ser265 instead of Phe/Tyr of the known FRP-interacting OCPs (Supplementary Fig. 12) [41].

## Discussion

This study was fueled by successful crystallization and determination of structures of OCP2 members from *C. epipsammum* PCC 9333 and *Gloeocapsa* sp. PCC 7428, representing cyanobacteria with completely different ecological, morphological and phylogenetic parameters. Prior knowledge of the OCP2 family was based mainly on few studies devoted nearly exclusively to TolyOCP2 [28,29,32]. At first glance, the availability of multiple OCP1 and recently obtained OCP3 structures [30,31] (Table 1) should have made homology modeling more robust and likely improved Alphafold performance on OCP, however, the final refined crystal structures differed significantly from the Alphafold models (Supplementary Fig. 5), especially in the functionally relevant regions ([30] and this work). These discrepancies and the fact that OCP is functional only when bound to a ketocarotenoid necessitated obtaining experimental structural data. Thus, the success with the OCP2 crystal structures completed the long-waited portfolio of crystal structures of all OCP families (Table 1), which enabled multiway comparison of their spectroscopic and functional properties, helped refine many cause-and-effect relationships and explain long accumulated observations.

Interestingly, members of different OCP families crystallize as monomers (OCP3) or a variety of dimers (OCP1, OCP2), yet all form stable monomeric species in solution, as validated by SEC-MALS of samples at as high as ∼0.5 mM protein concentration ([30] and this work). Purified OCP2 and OCP3 remain monomeric even upon drastic concentrating, which apparently excludes equilibrium association of monomers into dimers, questioning the stereotype that OCP2 is exclusively monomeric, whereas OCP1 and OCP3 are dimeric [28,29].

OCP1, OCP2 and OCP3 monomers retain the same overall structural organization, yet differ by the interdomain contacts, positions of the CTT and highly variable interdomain linkers, which all are strictly functionally relevant. Indeed, while the CTT was shown to play an important role in carotenoid uptake [49,50], the linker contributes to the kinetics of photoactivation and R-O relaxation [29]. The complete set of structures has allowed us for the first time to rationally correlate the properties of the absorbance spectra and protein-carotenoid interactions, which in particular explained the pronounced vibronic structure and the blue-shift of the OCP2 spectrum (coinciding with that of specific OCP1 mutants with only one effective H-bond donor to the carotenoid keto group) on one side and the broadened and red-shifted OCP3 spectrum on the other side. In the latter case, we identified the existence of a peculiar *chalcogen* bond (σ-hole interaction [55,56]) with the OCP3-specific Met251 residue and validated this by QM/MM calculations. PDB mining yielded many instances of analogous chalcogen bonds directly involved in protein-ligand interactions, including photoactive/photosensory proteins, where such tentatively important chalcogen bonds have not been explicitly analyzed and even mentioned (Supplementary Table 1). We show that in OCP3 and all-OCP ancestors [41], the chalcogen bond involving Met251 in addition to the two H-bonds between the carotenoid keto group and Tyr201 and Trp288 is accompanied by the broadening and red-shift of the absorbance spectrum. This finding indicates that the effect of Met residues should be more carefully analyzed in different systems in the future. For instance, one of the unreported chalcogen bonds is very probable in the recently published OCP3 structure from *N. flagelliforme* (PDB ID: 7YTF [31]), between the canthaxanthin’s ketogroup and Met125 in the NTD. The M125I mutation, obviously disrupting this bond, changes the spectrum and restores photoactivity, which was barely exhibited by this wild-type OCP3 [31]. We note that other Met residues of OCPs (e.g., Met202, Met205/206) can contribute to the complex interactions affecting the carotenoid anchoring in the CTD (Fig. 3B), and their role certainly warrants further investigations, despite the corresponding distances exceeding 4 A, a presumable threshold for such interactions [55]. Nevertheless, these interactions may play an important role in transition states.

Although crystal structures provide mainly static information and can select certain states, they capture all possible contacts of the carotenoid keto group with two (with Y and W, in OCP1 and OCP2) and three (with Y, W and M in OCP3) amino acid residues in the dark-adapted state. Combining spectroscopy with mutagenesis and structural data, we decomposed the spectra and assigned its components to OCP states with a different number of realized contacts (one, two or three). In particular, the possibility of OCP photoactivation is due to its state with only one bond being effectively present. States with a larger number of contacts reduce the vibronic spectral shape and likely decrease the OCP photoactivation probability, whereas states with no bonds characterize the photoactivated OCP^R^. This spectral and associated functional heterogeneity likely meets the objectives of specific OCP to balance photoactivation and relaxation rates for an optimal equilibrium of photoprotection and photosynthesis.

The complete portfolio of OCP structures also facilitated the analysis of the regulation by FRP. While previous studies stated that FRP does not bind to nor regulate OCP2 [28,29], we noticed the counterproductive binding of FRP to OCP2, which decelerated R-O transition at all temperatures tested, for both studied OCP2 proteins, albeit to a different degree. A similar phenomenon has recently been reported for SynFRP interaction with the ancestor of OCP [41]. Our analysis of the electrostatic potential distribution indicated that several signature residues on the NTD in OCP2 (and OCP3) reverse the charge of the NTD within the extended FRP-docking site, disturbing charge complementarity between the two proteins, which apparently disrupts the domain bridging activity of FRP [24]. The absence of FRP binding to OCP3 is likely due to its charge-reversed NTD and also slightly different CTD surface [30,41], since the isolated CTD of GbacOCP3 cannot recruit FRP, in contrast to the CTD of OCP1 (Supplementary Fig. 12 and [57]). The appearance of Ser265 in GbacOCP3 in place of Tyr/Phe present in OCP2, OCP1 and all-OCP ancestor likely accounts for the inefficiency of GbacOCP3/FRP interaction [41].

The fundamental ability of OCP1 and OCP3 to quench the model PBS from *Synechocystis* was comparable, although we observed the marked difference between CrinOCP2 and GcapOCP2, which differ by the residue-155. Structural analysis showed that, unlike Arg155 present in CrinOCP2 (OCP1 and OCP3), Gln155 present in GcapOCP2 cannot form authentic salt-bridges stabilizing the GcapOCP2-PBS and NTD-CTD interactions. Consistently, Gln155-containing TolyOCP2 was also a less efficient PBS quencher than TolyOCP1 [28].

This Gln155 feature is likely linked also with the slower R-O relaxation of GcapOCP2 than CrinOCP2. While the R-O transition of CrinOCP2 and GcapOCP2 was slightly faster than GbacOCP3 and much faster than SynOCP1, the two OCP2 species also differed substantially in their photocycling properties (Table 3), likely reflecting physiological adaptations of the corresponding OCP2-encoding cyanobacteria and the intermediate position of the OCP2 clade on the phylogenetic tree of the OCP superfamily. Clearly, more data on new OCP members should be obtained for making generalizations and we can anticipate exceptions from the established signatures of the OCP families. Indeed, the fast photoswitching dynamics is not an exclusive signature of OCP2 (and OCP3) as the recently described *Planktothrix* OCP1 is much faster than SynOCP1 and is still accelerated by FRP [33], in contrast to OCP2 and OCP3. Likewise, CrinOCP2 has an unusually low thermal optimum of photoactivation (35 °C), and unusual ability to be thermally activated in the dark (at >40 °C; Supplementary Fig. 8).

Our work may facilitate further analyses of various OCP representatives and inform the structure-guided design of novel OCP-based optogenetic and split molecular systems [35–38], and help engineer the OCP/FRP photoprotection system in biotechnologically relevant strains for improved cyanobacterial biomass production [39]. In plants, a fast-operating recovery from photoprotection indeed led to higher productivity [40].

## Methods

### Phylogenetic and bioinformatic analysis

Protein sequences of full-length OCP homologs (OCP1, OCP2 and OCP3 representatives) were retrieved from the dataset [29]. Our set was technically restricted to 150 sequences to match alignment requirements of T-COFFEE [58]. A tree was built using MEGA11 with the Maximum Likelihood method based on the JTT matrix-based model [59]. The analysis involved 69 OCP1, 23 OCP2 and 58 OCP3 amino acid sequences and considered 301 amino acid positions (gaps were eliminated). Initial tree(s) for the heuristic search were obtained automatically by applying Neighbor-Join and BioNJ algorithms to a matrix of pairwise distances estimated using a JTT model, and then selecting the topology with a superior log likelihood value. Family clades were determined by manual inspection and supported by a high percentage (>90%) of trees in which the associated taxa clustered together. Several OCP sequences cannot be reliably defined as representatives of OCP3 (=OCPX) clade according to our analysis and criterion, so for them we used the classification of [29]. The resulting tree is drawn to scale, with branch lengths measured in the number of substitutions per site.

### Proteins

cDNA corresponding to OCP2 from *Crinalium epipsammum* PCC 9333 (Uniprot K9VSN3) and *Gloeocapsa* sp. PCC 7428 (Uniprot K9XH36) were codon-optimized for expression in *E. coli*, synthesized by Synbio Technologies (China) and cloned into the pET28a+ vector (kanamycin resistance) using the *Nde*I and *Xho*I restriction sites. The plasmid design left extra N-terminal GSH… residues after cleavage by thrombin and included a glutamate-to-alanine mutation in position corresponding to Glu25 of *Gloeobacter* OCP3a, since this replacement promoted crystallization of OCP3a without interfering with its functional properties [30]. The individual CTD of *Gloeobacter* OCP3a was obtained in this work using *Pfu* DNA polymerase and the standard T7 reverse and a custom GlOCPX-CTD-NdeI-forw primer 5’-ATTATCATATGGTAGATGGAATCAATGATCC-3’ containing an *Nde*I site for cloning back into pET28-his-3C vector to express CTD in either apo– or ECH/CAN bound form as described earlier for CTD(SynOCP1) [24,57].

FRP (Uniprot P74103) and OCP1 (Uniprot P74102) from *Synechocystis* sp. PCC 6803 as well as OCP3a (formerly, OCPX) from *Gloeobacter kilaueensis* JS1 (Uniprot U5QHX0) were obtained in previous works as the electrophoretically homogeneous preparations devoid of tags [24,30]. The ΔNTE mutant lacking residues Met1-Phe12 from the N terminus of SynOCP1 was obtained earlier [24]. *Arthrospira maxima* FRP [Uniprot B5W3T4 (Limnospira (formerly, *Arthrospira*) *maxima* CS-328), coincides with Uniprot H1W9V5 (*Limnospira indica*) and K1X0E1 (*Arthrospira platensis* C1)] was recombinantly expressed and purified earlier [60]. In the final form, all proteins had His-tags cleaved. All constructs were verified by DNA sequencing in Evrogen (Moscow, Russia).

For obtaining the echinenone (ECH) bound protein forms, the OCP plasmids obtained were transformed into C41(DE3) *E. coli* cells carrying the pACCAR25ΔcrtXZcrtO plasmid (chloramphenicol resistance) encoding the gene cluster *crtY*, *crtI*, *crtB*, *crtE* and *crtO* sequences from *Erwinia uredovora*, which enabled ECH expression [30,47]. Protein apoforms were obtained in plain C41(DE3) cells. Protein expression was induced by adding 0.5 mM IPTG and continued for 48 h at 25 °C. Based on previous works [57,61], these conditions promoted the formation of ECH bound OCP forms.

Protein purification included two steps of immobilized metal-affinity chromatography separated by a thrombin-catalyzed His-tag cleavage and thrombin removal on a heparin column (GE Healthcare), hydrophobic interaction chromatography (HIC) on a 5-ml Phenyl-Sepharose column (GE Healthcare), and gel filtration on a Superdex 75 16/600 (GE Healthcare). Electrophoretically homogeneous preparations were obtained. Typical Vis/UV absorbance ratios for OCP2 samples obtained were in the 1.8-2.0 range as they were almost free from the apoprotein. Purified proteins were stored at –80 °C.

Phycobilisomes from *Synechocystis* were isolated in the present work as described earlier [62].

### Reverse phase high-performance liquid chromatography (HPLC)

The carotenoid content in the GcapOCP2 and CrinOCP2 preparations was assessed using RP-HPLC on a Nucleosil C18 4.6*250 column at a flow rate of 1 ml/min and column oven temperature of 28 °C. 100 μl of protein sample was mixed with 100 μl acetone and 100 μl hexane, then vortexed and centrifuged, and the colored supernatant containing carotenoids was dried under N_2_ atmosphere, then redissolved in 25 μl acetone, of which 20 μl were loaded onto the C18 column. Carotenoid standards (CAN, ECH and β-carotene) were used for reference. The elution profile was followed by absorbance at 460 nm and developed as follows: 5 min 70% acetone in water, 5-30 min gradient of 70->100% acetone in water, 30-32 min 100->70% acetone, then 70% acetone until 40 min.

### Spectrochromatography

To simultaneously record absorbance spectra and size-exclusion profiles of OCP species we used analytical size-exclusion spectrochromatography (ASESC). Protein samples (50 μl) were loaded on a Superdex 200 Increase 5/150 column (GE Healthcare, Chicago, Illinois, USA) pre-equilibrated with a 20 mM Tris-HCl buffer, pH 7.6, containing 150 mM NaCl (SEC buffer) and operated by a ProStar 210 system equipped with a diode-array detector Prostar 335 (Varian Inc., Melbourne, Australia). Flow rate was 0.45 ml/min. Absorbance in the 240-900 nm range was registered with 1-nm precision (4 nm slit width) and a 5 Hz frequency during each run. Each experiment was repeated at least three times, and the most typical results are presented. To assess Mw of the protein peaks obtained, the column was pre-calibrated using α-lactalbumin (15 kDa), bovine serum albumin monomer (66 kDa), bovine serum albumin dimer (132 kDa) and bovine serum albumin trimer (198 kDa).

### Interaction of OCP species with FRP

ASESC was also used to track the ability of OCP species (ΔNTE-SynOCP1 holoform, SynOCP1 WT apoform, or the apoforms of CrinOCP2 and GcapOCP2) to form a complex with SynFRP. To this end, 10 μM of OCP and 20 μM of FRP were pre-incubated in SEC buffer for 15 min at room temperature and then loaded onto a Superdex 200 Increase 5/150 column (GE Healthcare) pre-equilibrated with the same buffer at 0.45 ml/min, while the elution profiles were followed by either 280 or 460 (or 500) nm absorbance.

The structure of the 1:2 OCP:SynFRP complex was predicted by ColabFold [63] for various OCP variants using sequences of an OCP monomer and SynFRP dimer, and the models were relaxed after prediction runs to minimize clashes. To analyze the effect of SynFRP on various OCP we applied the APBS plug-in for PyMol (using default parameters) to depict surface electrostatic potential distribution, colored from positive (+3 k T e^−1^; blue) to negative (−3 k T e^−1^; red).

### Absorbance measurements of the OCP photocycle

To monitor the photocycle of various OCP forms (10 μM) in the absence or the presence of SynFRP (20 μM), protein samples (100 μl) in Tris-HCl buffer pH 7.6 containing 150 mM NaCl were transferred to a 1 cm quartz cuvette and the steady-state absorbance spectrum was continuously recorded. A blue light-emitting diode (M455L3, Thorlabs, USA) with a maximum emission at 445 nm was used to photoswitch OCP via continuous illumination until reaching the equilibrium corresponding to the specific temperature, which was maintained by a Peltier-controlled cuvette holder Qpod 2e (Quantum Northwest, USA). After reaching the equilibrium, the blue LED was switched off and the R-O transitions were followed by changes of absorbance at 550 nm until reaching a plateau. Each experiment was repeated at least three times, and the most typical results are presented. For data presentation and comparison we normalized the kinetic R-O curves obtained under different conditions (a combination of temperature, OCP variant and the presence or absence of SynFRP) to 0,100 and then used the logarithm of the X axis (time) to facilitate comparison of the very different time scales. Time required to convert 50% of the light-adapted form back to the dark-adapted form (τ_1/2_) was used to compare different rates of the R-O transition. In addition, for GcapOCP2 we build an Arrhenius plot. To this end, kinetics of the R-O transition in the temperature range 5-23 C were approximated by 2 exponential decay and the corresponding mean time constant t_m_ for each temperature was derived. Ln(1/t_m_) (rate constant) against 1000/T (K) was linear and was used to derive Ea as described earlier [30,64].

To observe the stages of the photocycle on an ms time scale, OCP photoactivation was initiated by 15-100 ms flashes of high-power LED (up to 5 W) SOLIS-445C (Thorlabs, USA) operating under the control of DC2200 (Thorlabs, USA), generating actinic flashes with duration ranging from 3 μs to 10 s. A stabilized deuterium lamp SL S204 (Thorlabs, USA) was used to probe the change in absorbance of the photoactivated OCP sample. A FLAME + spectrometer (Ocean Insight, USA), with a diffraction grating providing a spectral resolution of 10 nm, was used to record changes in the optical density of the sample with 2 ms time resolution. Changes in optical density were recorded at 550 nm to observe changes in the concentration of red forms of OCP, as well as at 445 nm to determine the moment of actinic flash on and off. The temperature of the sample was controlled by a Qpod 2e (Quantum Northwest, USA) cuvette holder. To improve the signal-to-noise ratio, each experiment was repeated at least 30 times.

### Time-resolved PBS fluorescence measurements

PBs fluorescence decay kinetics in 0.75 M Na-phosphate pH 7.0 containing 0.75 M sucrose with picosecond time resolution were recorded with a time– and wavelength-correlated single photon counting setup based on an HMP-100-40 detector and an SPC-150 module (Becker&Hickl, Germany). Fluorescence was excited at 620 nm by 150 fs flashes generated by optical parametric generator TOPOL (Avesta Project Ltd., Russia) PBs emission was recorded at 680 nm. Fluorescence decay was approximated by a sum of two exponential decay functions with the SPCImage (Becker&Hickl, Germany) software package, considering the incomplete decay of the states with long lifetimes at 80 MHz repetition rate. The temperature of the sample was controlled by a Qpod 2e (Quantum Northwest, USA) cuvette holder. Photoactivation of OCP was triggered by a blue LED (445 nm, 200 mW, Thorlabs, USA).

### SEC-MALS

Size-exclusion chromatography coupled to multi-angle light scattering (SEC-MALS) of GcapOCP2 or CrinOCP2 holoform samples (1 mg/ml in 100 μl) was carried out on a Superdex 200 Increase 10/300 column (GE Healthcare) connected to a tandem of consecutive detectors Prostar 335 (Varian, Australia) and a miniDAWN (Wyatt Technology, USA). The column was equilibrated with filtered (0.1 μm) and degassed 20 mM Tris-HCl buffer, pH 7.6, containing 150 mM NaCl, and was operated at 0.8 ml/min. Data were processed in ASTRA 8.0 (Wyatt Technology, USA) using dn/dc = 0.185 and protein extinction coefficients (mg/ml^-1^ cm^-1^) at 280 nm equal to 1.52 (GcapOCP2(ECH)) and 1.50 (CrinOCP2(ECH)). Such extinction coefficients were calculated using the experimental absorbance spectra of OCP2(ECH) complexes and assuming that ECH absorbance at 280 nm is ∼⅛ of its absorbance at 500 nm. To account for subtle 660-nm laser absorbance (less than 2%), the laser intensity was corrected by forward laser power.

### Crystallization of GcapOCP2 and CrinOCP2

An initial crystallization screening of untagged GcapOCP2 and CrinOCP2 was performed with a robotic crystallization system (Oryx4, Douglas Instruments, UK) and commercially available crystallization screens (Hampton Research, USA) using sitting drop vapor diffusion at 15 °C. Protein samples (12.6 mg/ml GcapOCP2 and 11.5 mg/ml CrinOCP2) were prepared in 20 mM Tris-HCl buffer, pH 7.6, 150 mM NaCl, 3 mM NaN_3_. The drop volumes were 0.2 µl with 1:1 protein-to-precipitant ratio and 0.25 µl with 3:2 protein-to-precipitant ratio, respectively. The plates were covered by aluminum foil during crystallization. While CrinOCP2 readily crystallized in different chemical conditions, only one crystal of GcapOCP2 could be obtained, which grew during 1 month. Crystals suitable for X-ray analysis were obtained in the following conditions: 0.2 M Magnesium formate dihydrate, 0.1 M Sodium acetate trihydrate pH 4.0, 18% w/v PEG MME 5000 (GcapOCP2) and 0.1 M Imidazole pH 6.5, 1.0 M Sodium acetate trihydrate (CrinOCP2). Of note, our previously determined crystal structures of SynOCP1 revealed no significant conformational variation in the pH range from 4 to 7 [19], suggesting that the conformation of also GcapOCP2 is representative of the native state.

### X-ray data collection, structure determination and refinement

Crystals were briefly soaked in cryo-solution containing 20% glycerol (Hampton Research, USA) immediately prior to diffraction data collection and flash-frozen in liquid nitrogen. The data were collected at 100K using BL17B1 beamline (SSRF, China) and Rigaku OD XtaLAB Synergy-S home source (Moscow, Russia) for GcapOCP2 and CrinOCP2, respectively. Indexing, integration and scaling were done using XDS [65], CrysAlisPRO (Oxford Diffraction / Agilent Technologies UK Ltd, Yarnton, England) and Aimless [66] (Table 2).

Both structures were solved by molecular replacement with Phaser [67] using the structure of OCP1 from *Synechocystis* sp. PCC 6803 (PDB ID: 5TUX) and an AlphaFold model as search models for GcapOCP2 and CrinOCP2, respectively. The refinement was carried out using REFMAC5 [68]. The isotropic individual atom B-factors as well as hydrogens in riding positions were used during the refinement. The visual inspection of electron density maps and manual model rebuilding were carried out with COOT [69].

### Molecular modeling

Non-H atom coordinates of GbacOCP3 (PDB ID 8A0H [30]) were used to set-up calculations. Hydrogen atoms were added assuming deprotonated side chains of aspartate and glutamate residues, positively charged side chains of lysine and arginine residues; other residues were neutral. The macromolecule was described using CHARMM36 [70,71], water molecules with TIP3P [72] and carotenoid with CGenFF [70] force field parameters. The system was solvated in a rectangular water box with the total size of the system being 42,832 atoms. Preliminary equilibration was performed via classical force field parameters in molecular dynamics simulations in the canonical NPT ensemble at 300K and 1 atm. 10 ps QM/MM MD equilibration and subsequent 10 ps production run was performed to analyze hydrogen bond flexibility in the keto group region of the carotenoid. The representative frame was utilized to obtain a minimum on the potential energy surface. It was utilized to perform topological analysis according to the QTAIM theory [73] and quantification of the electron density at bond critical points corresponding to the interactions of the oxygen atom of the carotenoid. The QM subsystem included a fragment of the carotenoid and side chains of neighboring residues, Y202, M251 and W289 (Fig. 3E, ii). The QM subsystem was treated at the Kohn-Sham PBE0-D3/6-31G** level [74,75]. Vertical electronic excitation energy was calculated at the CAM-B3LYP/def2-SVP level [76]. NAMD2 [77], TeraChem [78], ORCA [79] and Multiwfn [80] programs were utilized for computations and VMD [81] for visualization.

## Data availability

Atomic coordinates and structure factors have been deposited in the Protein Data Bank under the following accession codes: 8PYH (CrinOCP2) and 8PZK (GcapOCP2). All other data associated with the study are available upon request.

## Abbreviations used

ASESC: analytical size-exclusion chromatography
CAN: canthaxanthin
ECH: echinenone
hECH: 3-hydroxyechinenone
CrinOCP2: Crinalium epipsammum OCP clade 2
CTD: C-terminal domain
CTDH: C-terminal domain homolog
DA: dark-adapted
FRP: fluorescence recovery protein
GcapOCP2: Gloeocapsa sp. PCC 7428 OCP clade 2
OCP: Orange Carotenoid Protein
GbacOCP3: *Gloeobacter kilaueensis* OCP clade 3
HCP: helical carotenoid protein
IMAC: immobilized metal-affinity chromatography
IPTG: isopropyl-β-thiogalactoside
LA: light-adapted
LED: light-emitting diode
NPQ: nonphotochemical quenching
NTD: N-terminal domain
NTE: N-terminal extension
PBS: phycobilisomes
SDS-PAGE: sodium dodecyl sulfate-polyacrylamide gel electrophoresis
SEC: size-exclusion chromatography
SEC-MALS: size-exclusion chromatography coupled to multi-angle light scattering
SynOCP1: *Synechocystis* sp. PCC 6803 OCP clade 1.

## Acknowledgments

N.N.S. is thankful to Dr. Yaroslav Faletrov for help in finding examples of the chalcogen bonds in PDB and for fruitful discussions, to Andrei O. Zupnik for providing CTD(GbacOCP3) preparations and to Dr. Mikhail E. Minyaev for access to the Rigaku OD XtaLAB Synergy-S diffractometer. The study was supported by the Ministry of Science and Higher education of the Russian Federation in the framework of the Agreement no. 075-15-2021-1354 (07.10.2021). Spectroscopic characterization of OCP photocycling transitions and OCP-PBS interactions was supported by the joint Russian Science Foundation/National Scientific Fund of China (grant number 21-44-00005 and 42061134020). SEC-MALS and crystallization screening were done at the Shared-Access Equipment Centre “Industrial Biotechnology’’ of the Federal Research Center “Fundamentals of Biotechnology” of the Russian Academy of Sciences. MGK acknowledges the use of supercomputer resources of the M.V. Lomonosov Moscow State University and of the Joint Supercomputer Center of the Russian Academy of Sciences.

## Author contributions

NNS – initiated and coordinated the study; YBS, NNS – expressed and purified proteins; LAV – crystallized proteins; YBS, EGM, NNS – designed and performed experiments; NAE – performed HPLC analysis; NNS, EGM, GVT, AYB – collected photoswitching kinetics data; LAV, KMB – collected X-ray data; NNS – solved structures; NNS, KMB – refined structures; MGK – performed molecular modeling; YBS, KMB, BG, SQ, EGM, NNS – analyzed data and discussed the results; NNS wrote the paper with input from EGM; VOP – supervised the study and acquired funding.

## Supplementary Tables

**Supplementary Table 1.** Exemplary protein-ligand complexed with the chalcogen bonds within 4 A distance between methionine’s sulfur and ligand’s oxygen atoms.

| Protein name | Ligand name | PDB ID | Met involved | S...O interatom distance | ref. |
| --- | --- | --- | --- | --- | --- |
| Insulin-like growth factor 1 receptor kinase domain | hydantoin | 3O23 | Met1079 | 3.0 Å | [84] |
| Phosphatidylinositol-4,5-bisphosphate 3-kinase catalytic subunit, gamma isoform | AS-604850 inhibitor | 2A4Z | Met953 | 3.0 Å | [85] |
| MAPKAP Kinase-2 | squarate Inhibitor | 3FPM | Met138 | 3.5 Å | [86] |
| Androgen receptor | bulky-B-ring antagonist | 3RLJ | Met895 | 3.5 Å | [87] |
| OCP3b (OCPX2) from *Nostoc flagelliforme* | canthaxantin | 7YTF | Met251 | 3.5 Å | [31] |
| OCP3b (OCPX2) from *Nostoc flagelliforme* | canthaxantin | 7YTF | Met125 | 3.6 Å | [31] |
| Estrogen Receptor | benzopyran | 2POG | Met343 | 3.7 Å | [88] |
| Estrogen Receptor | oxabicyclic derivative compound | 2QR9 | Met343 | 3.7 Å | [89] |
| OCP3a from *Gloeobacter kilaueensis* JS1 | echinenone | 8A0H | Met251 | 3.7 Å | [30] |
| Light–oxygen–voltage (LOV) fluorescent protein domain | flavin mononucleotide | 3SW1 | Met73 | 3.8 Å | [90] |
| Ketosteroid isomerase | equilenin | 1CQS | Met90 | 3.8 Å | [91] |
| Androgen receptor | bulky-B-ring antagonist | 3RLJ | Met742 | 3.9 Å | [87] |
| Xanthorhodopsin | salinixanthin | 3DDL | Met211 | 3.9 Å | [92] |
| Iron/alpha-ketoglutarate-dependent dioxygenase asqJ | 2-oxoglutaric acid | 6EOZ | Met122 | 3.9 Å | [93] |

## Supplementary Figures

**Supplementary Fig. 1.**
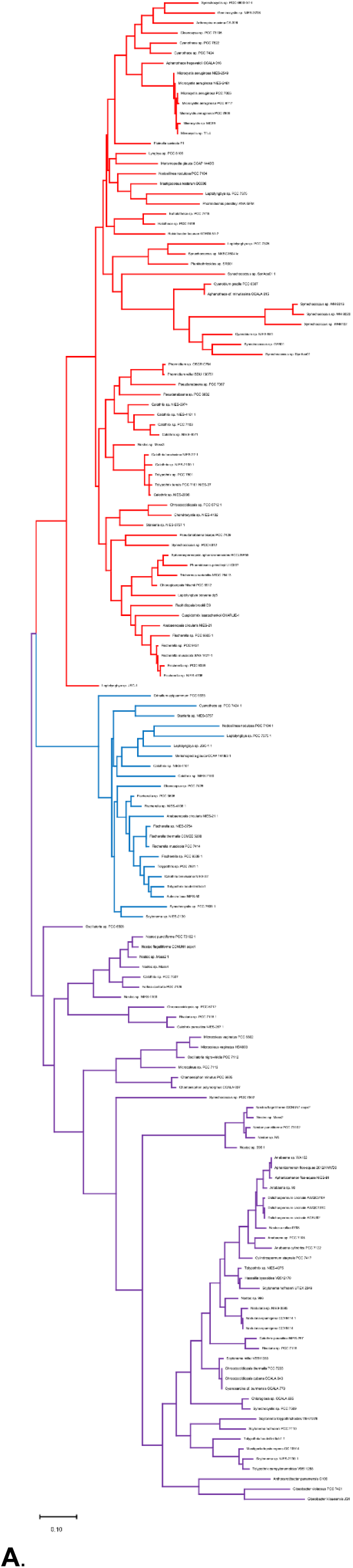
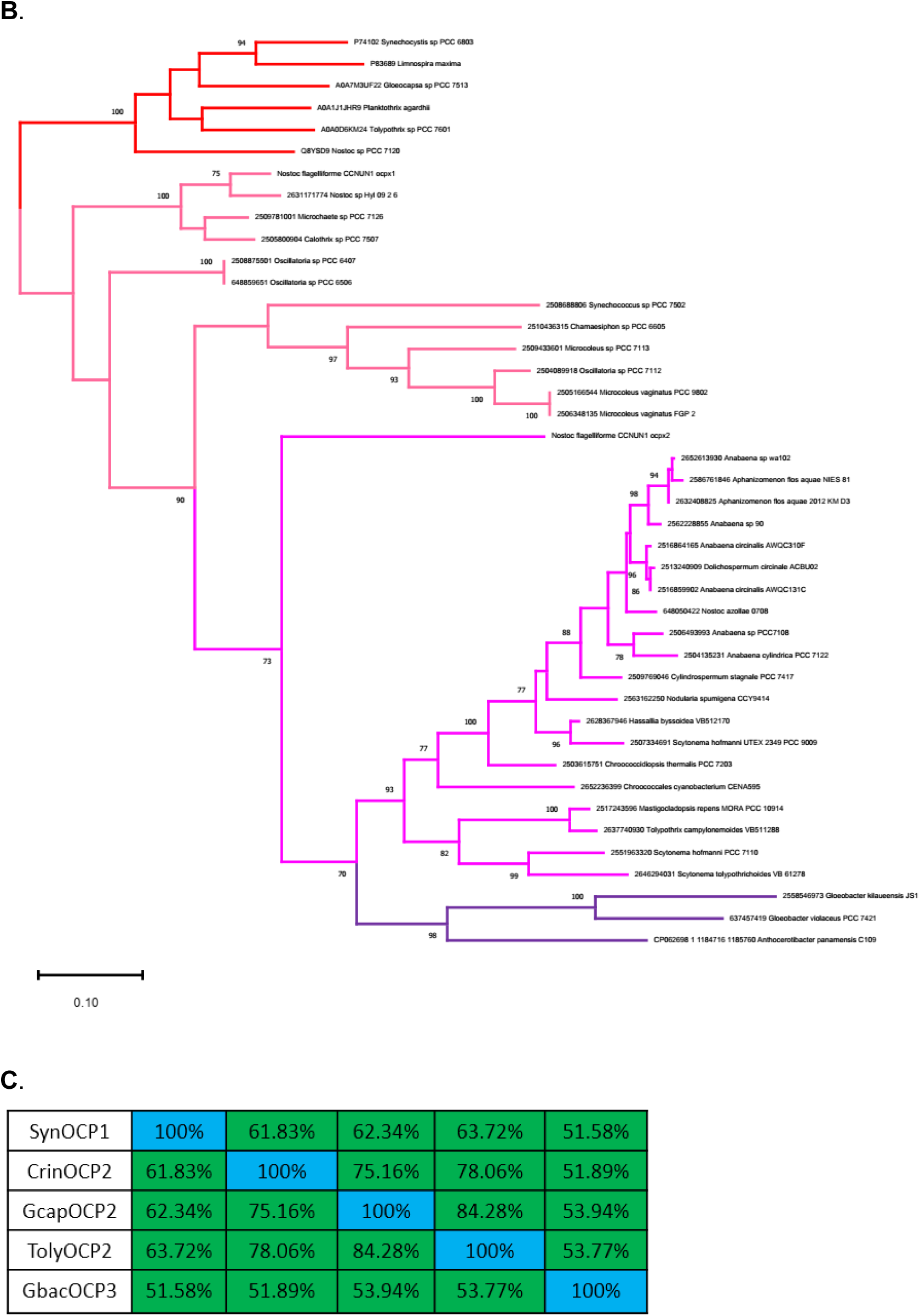
**A**. Complete phylogenetic tree of OCP families (shown in Fig. 1A in a shrinked form). OCP1 (red), OCP2 (blue), OCP3 (violet). **B**. Phylogenetic tree of OCP3 family showing the subdivision to OCP3a (violet), OCP3b (magenta) and OCP3c (pink), according to [30]. Selected OCP1 sequences were used as an outgroup (red). Note that OCPX2 [31] is subjectively classified as OCP3b, in fact OCPX2 being in between OCP3b and OCP3c subclades. **C**. Percent identity matrix for the sequences aligned in Fig. 1B.

**Supplementary Fig. 2.**
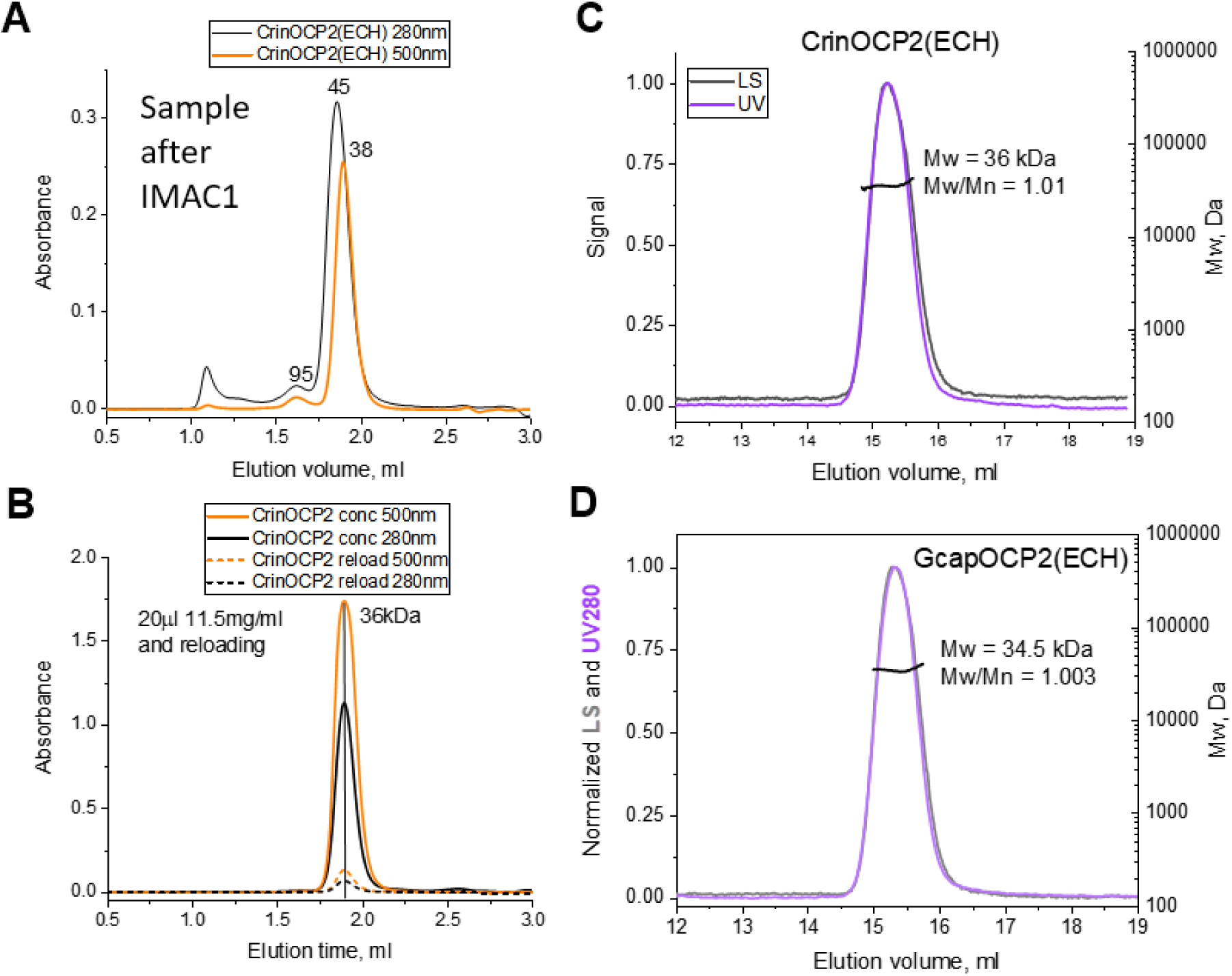
Analysis of CrinOCP2 oligomerization by SEC. **A,B**. ASESC of CrinOCP2 expressed in echinenone-producing *E.coli* cells, immediately after IMAC1 and after complete purification (Superdex 200 Increase 5/150 column (Cytiva); 0.45 ml/min). The CrinOCP2 peak collected during the run was reloaded on the column (∼10 times dilution) to study the effect of dilution. The apparent Mw for the peaks are indicated as determined from column calibration. **C,D**. Determination of the absolute Mw of the final CrinOCP2(ECH) (**C**) or GcapOCP2(ECH) (**D**) preparation using SEC-MALS (Superdex 200 Increase 10/300 column (Cytiva); 0.8 ml/min). Average Mw across the peak is indicated along with the polydispersity index (Mw/Mn).

**Supplementary Fig. 3.**
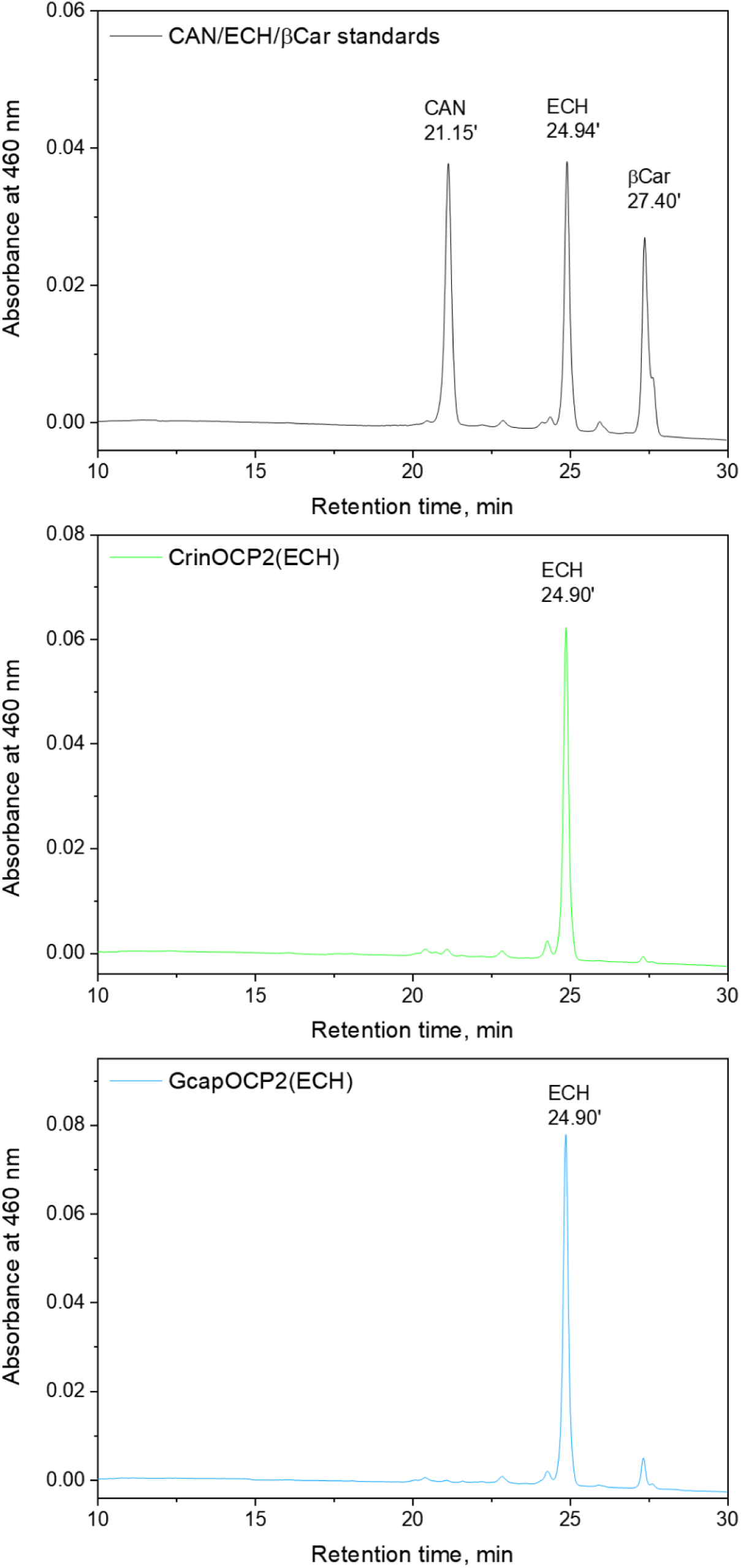
HPLC analysis of the carotenoids bound to CrinOCP2 and GcapOCP2. Temperature was 28 °C, flow rate 1.0 ml/min, Nucleosil C18 4.6*250 mm column, the gradient structure is described in Methods.

**Supplementary Fig. 4.**
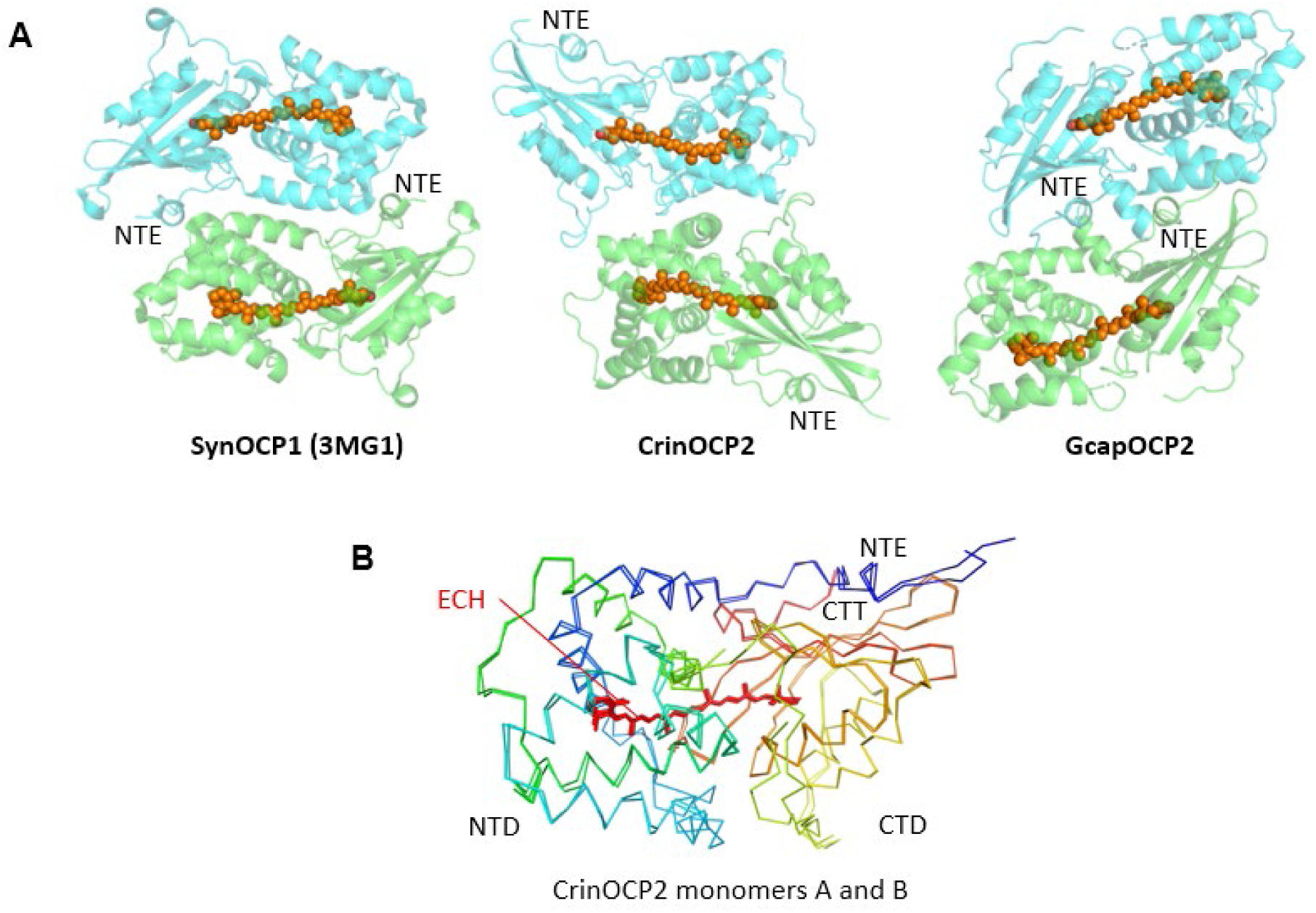
OCPs crystallize as different dimeric forms. **A**. Antiparallel crystallographic dimers of SynOCP1, CrinOCP2 and GcapOCP2 shown as ribbon diagrams with subunits colored by green and cyan. Echinenone molecules are shown by orange spheres. The positions of the N-terminal extensions (NTE) are marked for convenience. Note that the asymmetric units (ASU) of the crystals contained either dimer (SynOCP1 and CrinOCP2) or monomer (GcapOCP2), which formed the dimer with the symmetry-related monomer from neighboring ASU. **B**. Superposition of the two chains of CrinOCP2 shows the similarity of their conformation. The chains are colored by a gradient from blue (N terminus) to red (C terminus).

**Supplementary Fig. 5.**
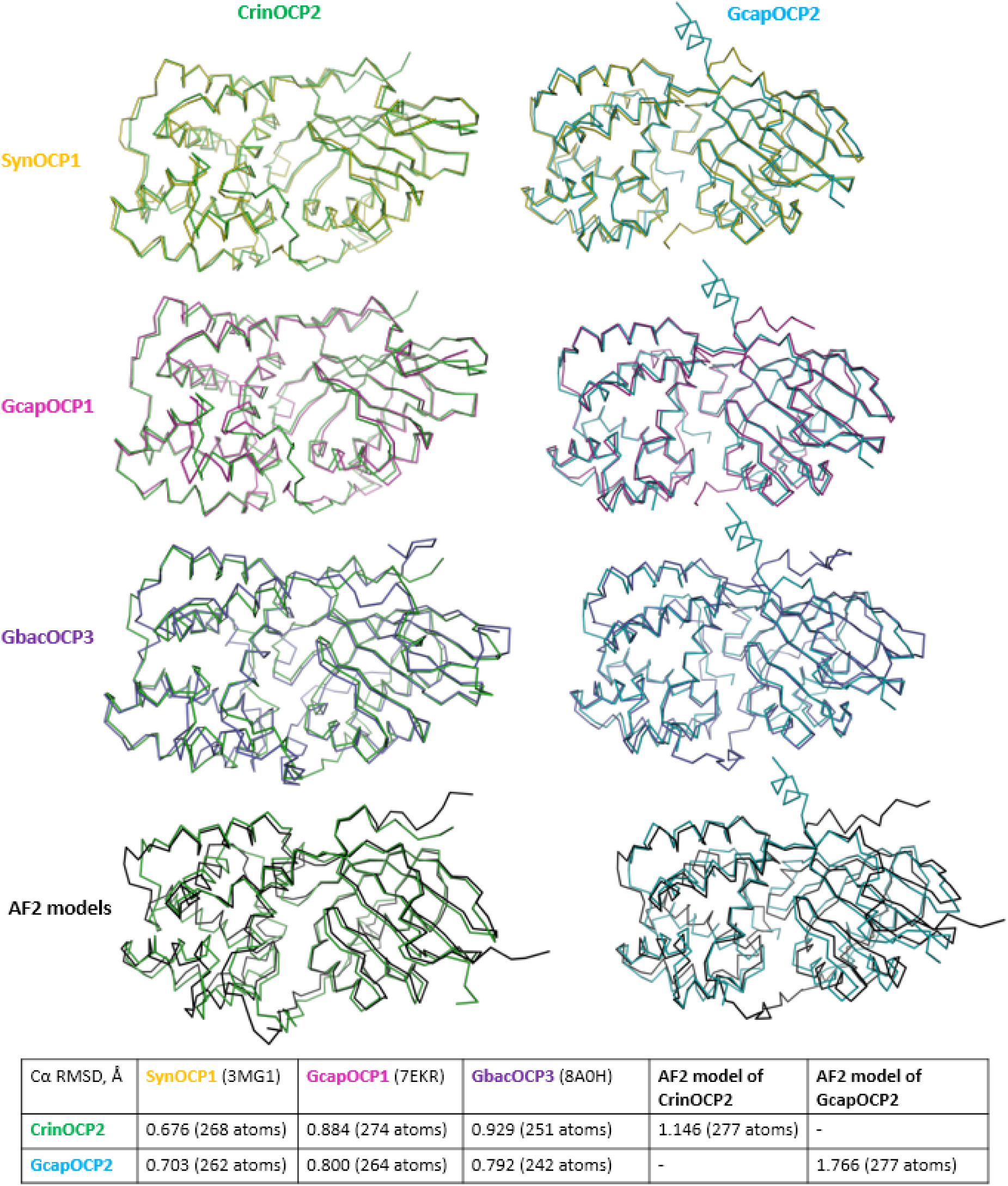
Superposition of the CrinOCP2 or GcapOCP2 crystal structures onto the structures of SynOCP1, GcapOCP1, GbacOCP3 or Alphafold models of CrinOCP2 or GcapOCP2. On the top, the pairwise backbone structural overlay for the corresponding OCP monomers are presented using the uniform color coding. On the bottom, the table for the corresponding Cα RMSD values upon structure superposition is shown. PDB identifiers are shown in parentheses.

**Supplementary Fig. 6.**
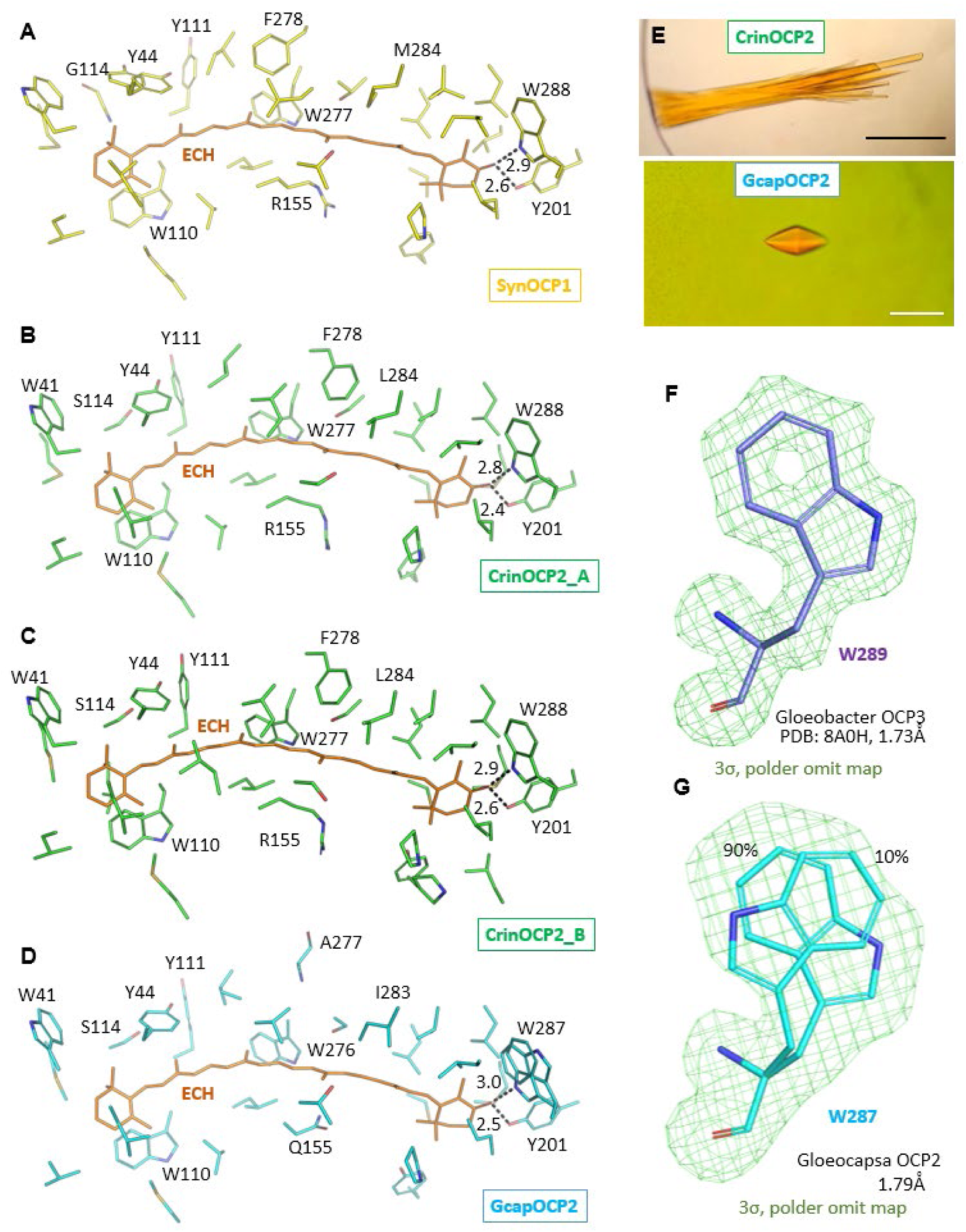
Carotenoid environment in OCP structures. **A-D**. Identical views of the echinenone binding sites in SynOCP1 (**A**), CrinOCP2 (two chains of the ASU are shown in **B** and **C**), and GcapOCP2 (**D**). Echinenone (ECH) is shown by orange sticks, protein chains are uniformly color coded as in the rest of the paper. Characteristic H bond lengths with the keto group of ECH are indicated in angstrom. **E**. CrinOCP2(ECH) and GcapOCP2(ECH) crystals. Scale bars correspond to 70 (black) and 20 (white) μm. **F,G**. Polder omit maps [94] for the key ECH contacting Trp residue in GbacOCP3 (**F**) and in GcapOCP2 (**G**) structures refined at a similar resolution, showing that only in GcapOCP2 the Trp side chain adopts two alternative conformations.

**Supplementary Fig. 7.**
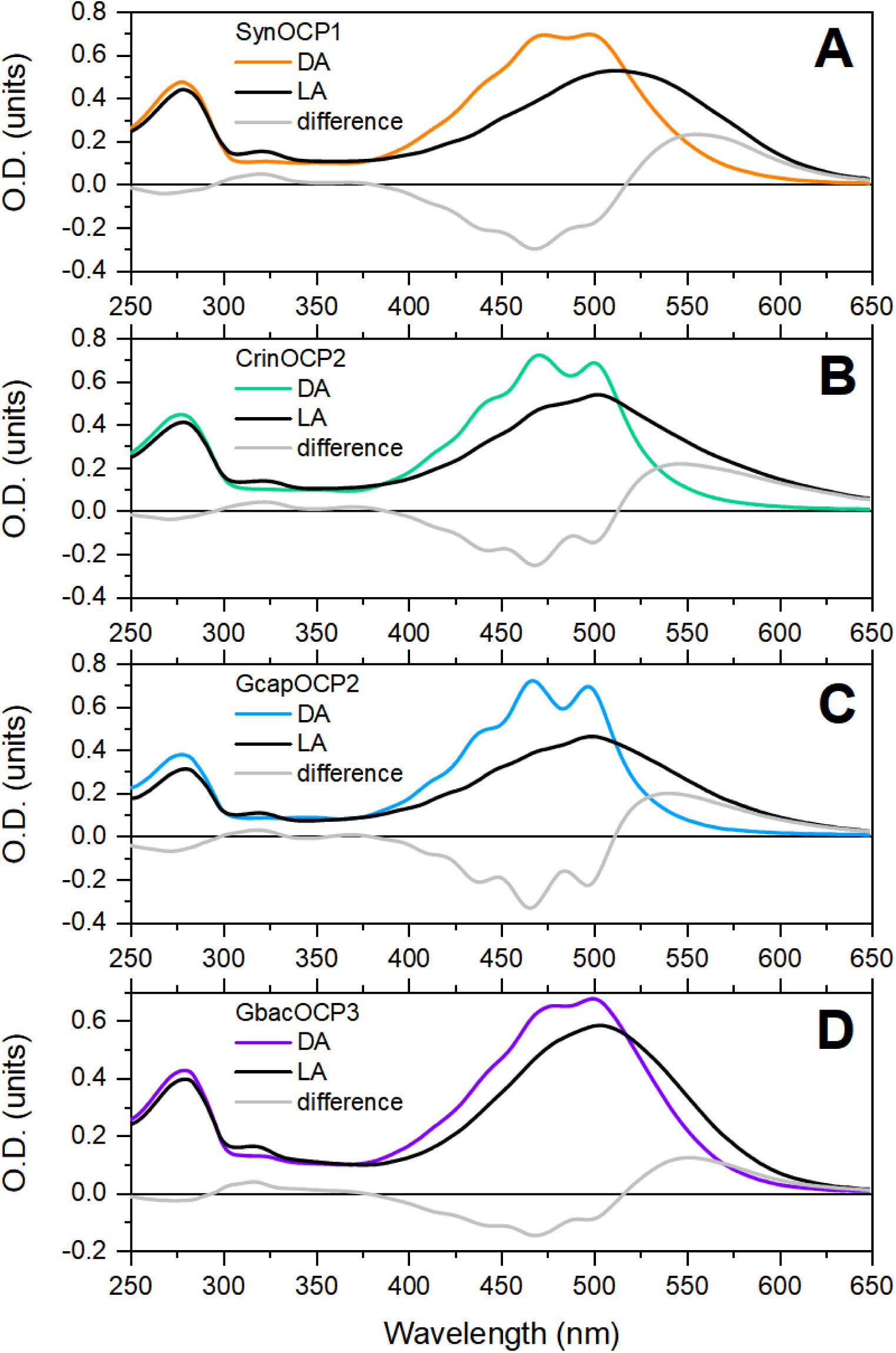
The absorbance spectra of SynOCP1 (**A**), CrinOCP2 (**B**), GcapOCP2 (**C**) and GbacOCP3 (**D**) in the dark-adapted (DA) and light-adapted (LA) states. Grey spectra represent LA minus DA spectra. All proteins were ECH-bound. Spectra were measured at 5 °C, for photoactivation proteins were irradiated with actinic light (445 nm, 2 W) for 120 seconds.

**Supplementary Fig. 8.**
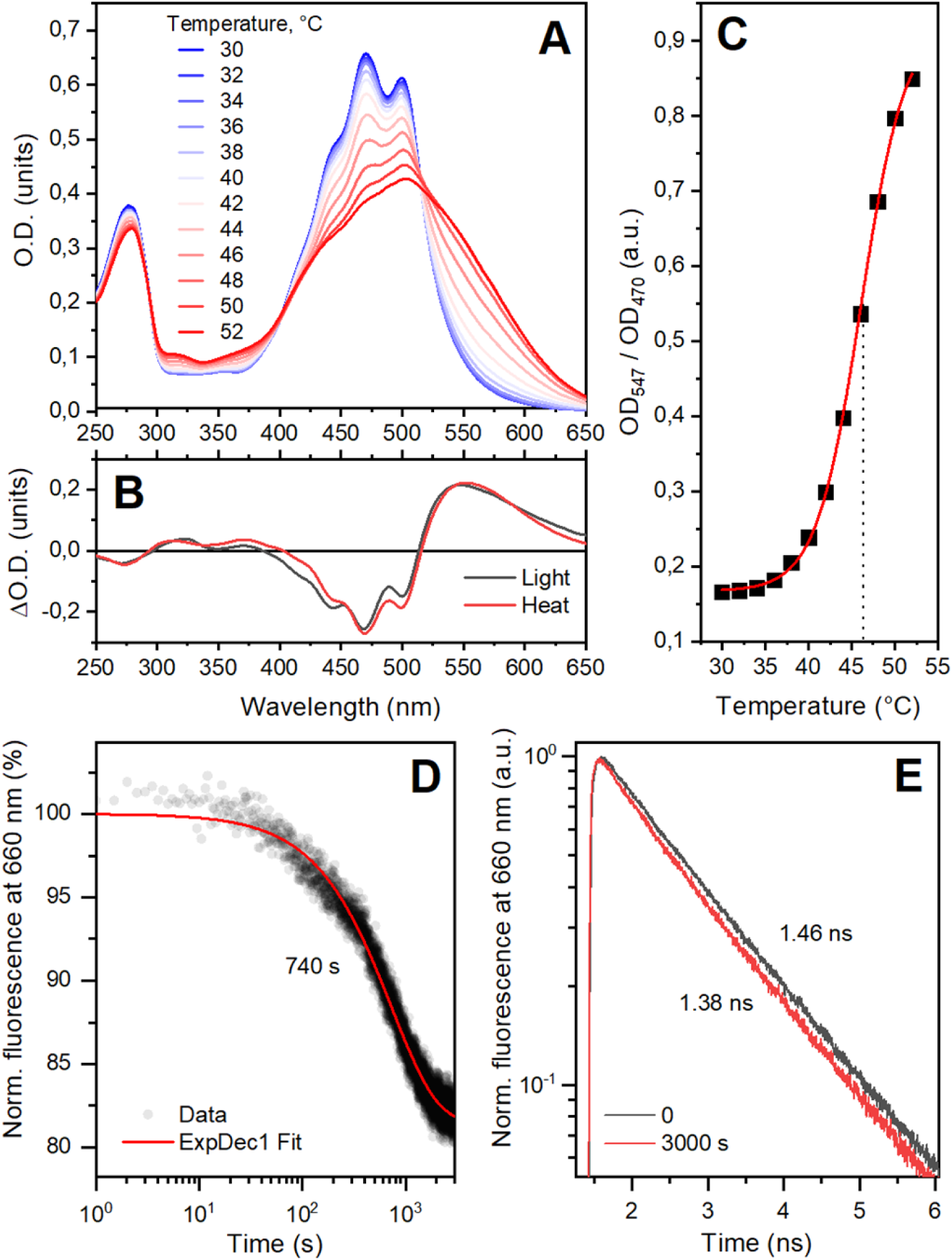
Temperature-induced activation of CrinOCP2. **A** – absorption spectra of CrinOCP2 at increasing temperature from 30 to 52 °C. **B** – comparison of difference absorption spectra of CrinOCP2 under “light minus dark” and “52 minus 30 °C” conditions. **C** – relative contribution of the red form to the absorption spectrum of CrinOCP2 at different temperatures calculated as the ratio of absorbance at 547 and 470 nm. **D** – change in fluorescence intensity of *Synechocystis* PBS upon incubation in the presence of the 100-fold excess CrinOCP2 at 45 °C. **E** – fluorescence decay kinetics of PBS fluorescence before the addition of CrinOCP2 and after 3000 s incubation of the PBS-OCP mixture at 45 °C. Numbers mark the characteristic lifetimes of the states. Fluorescence of PBS was excited at 590 nm.

**Supplementary Fig. 9.**
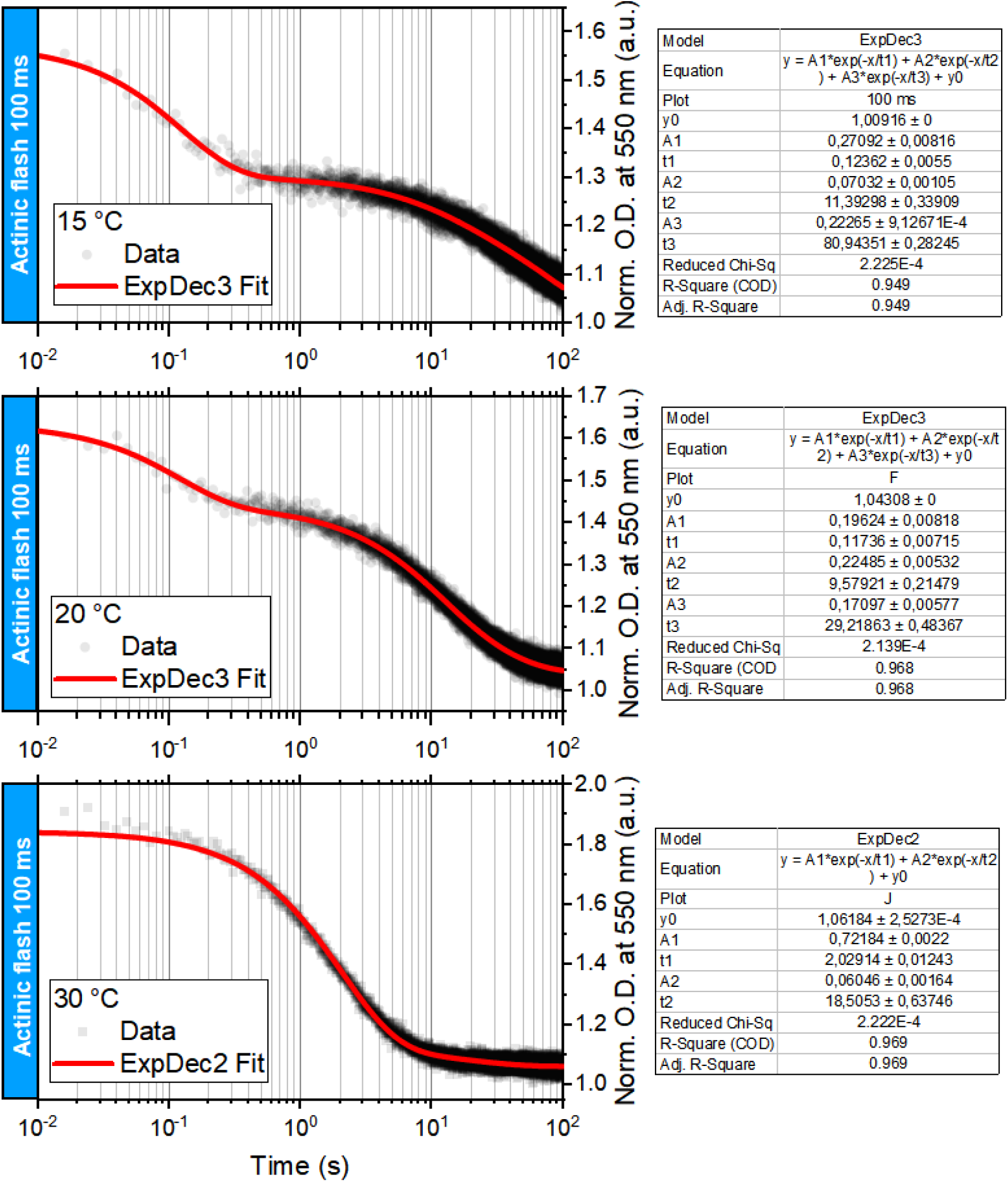
Examples of red form relaxation kinetics of GcapOCP2 at 15, 20, and 30 °C after a 100 ms flash of actinic light. The tables on the right show the parameters of the approximation of the data by the sum of decaying exponentials. Note that a simplified model with two time constants was used for 30 °C because the contribution of the fast component becomes negligible.

**Supplementary Fig. 10.**
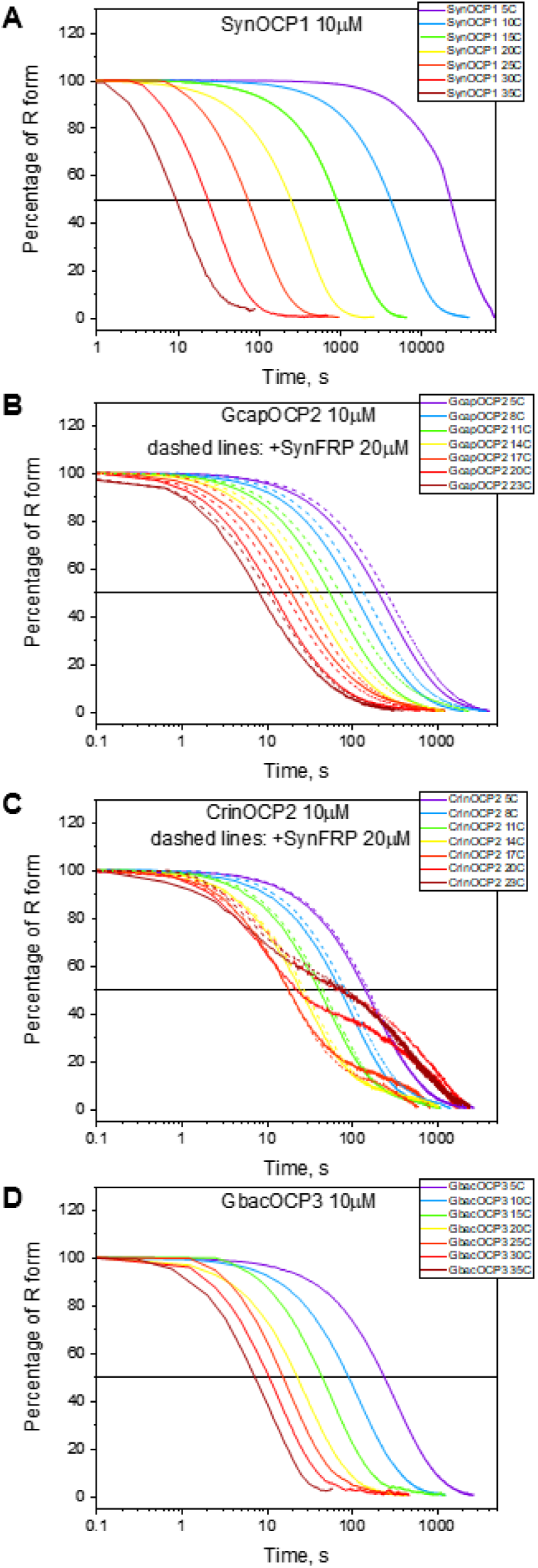
Temperature dependences of the R-O transition for various OCPs recorded after switching off blue LED illumination continued until reaching the equilibrium. Normalized temperature dependencies of the R-O transition curves for SynOCP1 (**A**), GcapOCP2 (**B**), CrinOCP2 (**C**) and GbacOCP3 (**D**) presented in semi-logarithmic coordinates. The R-O kinetic curves in the presence of SynFRP are shown for OCP2 proteins only (dashed lines on B and C, respectively). Zero time point corresponds to switching actinic light off. Concentrations of OCP and SynFRP were 10 and 20 μM, respectively, in 20 mM Tris-HCl pH 7.6, 150 mM NaCl buffer. In all cases, OCPs were ECH-bound.

**Supplementary Fig. 11.**
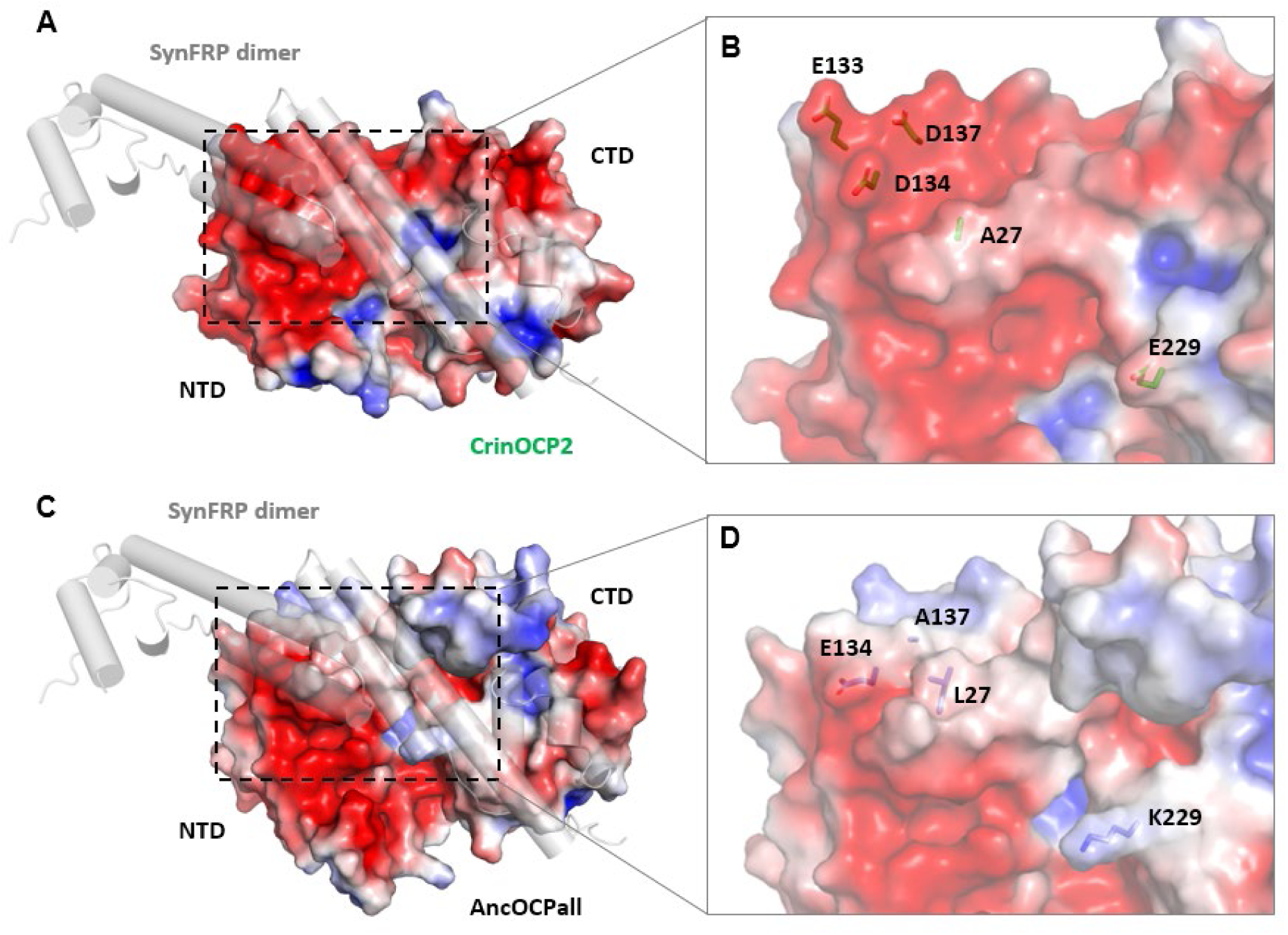
The electrostatic potential distribution on the surface of CrinOCP2 (**A,B**) and the ancestor of all OCP (AncOCPall, **C,D**) calculated by APBS tools [color gradient from red (–3 kT/e) to blue (+3 kT/e)] at 150 mM salt concentration. The crystal structure of CrinOCP2, chain B was overlaid onto the SynOCP1 chain of the SynOCP1/SynFRP dimer complex predicted by Alphafold, to illustrate the OCP1 dissimilar distribution of the electrostatic potentials within the FRP docking site (dashed rectangle). AncOCPall monomer structure was predicted by Alphafold using the amino acid sequence reconstructed in [41] and then overlaid onto the SynOCP1 chain of the Alphafold model of the SynOCP1/SynFRP dimer complex, similar to the procedure described for CrinOCP2. **A,C**. Overall views. **B,D**. Close-up views. Key residues at the FRP docking site are shown as sticks under the semi-transparent surface colored according to the electrostatic potential (scale as above).

**Supplementary Fig. 12.**
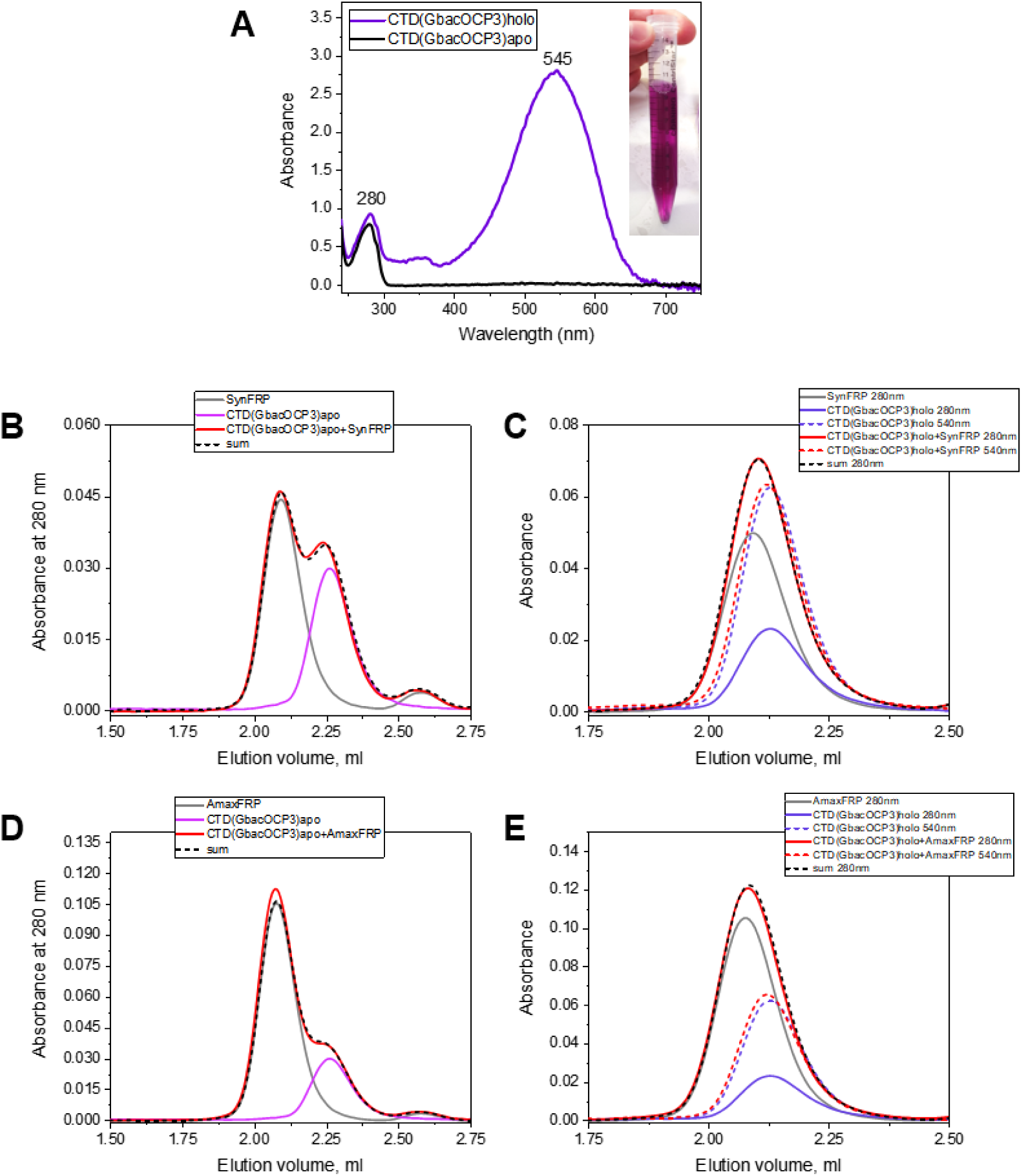
The isolated CTD of GbacOCP3 does not interact with either SynFRP or AmaxFRP. **A**. Absorbance spectra of the apo– or carotenoid bound CTD(GbacOCP3) recorded using a nanophotometer NP80 (Implen, Germany). The main peak maxima are indicated in nm. The insert shows the color of the holoprotein sample obtained using a combination of the subtractive IMAC and gel filtration as described earlier [57,95]. **B-E**. Analysis of the interaction between either *Synechocystis* FRP (**B,C**) or *Arthrospira maxima* FRP (**D,E**) and either the CTD(GbacOCP3) apoform (**B,D**) or the holoform (**C,E**). No significant difference is found for the FRP+CTD mixtures compared with the algebraic sums of the individual FRP and CTD profiles, indicating no stable association between these proteins.

